# Deletion of Neuroligins from Astrocytes Does Not Detectably Alter Synapse Numbers or Astrocyte Cytoarchitecture by Maturity

**DOI:** 10.1101/2023.04.10.536254

**Authors:** Samantha R. Golf, Justin H. Trotter, Jinzhao Wang, George Nakahara, Xiao Han, Marius Wernig, Thomas C. Südhof

## Abstract

Astrocytes perform multifarious roles in the formation, regulation, and function of synapses in the brain, but the mechanisms involved are incompletely understood. Interestingly, astrocytes abundantly express neuroligins, postsynaptic adhesion molecules that function as synaptic organizers by binding to presynaptic neurexins. Here we examined the function of neuroligins in astrocytes with a rigorous genetic approach that uses the conditional deletion of all major neuroligins (*Nlgn1-3*) in astrocytes *in vivo* and complemented this approach by a genetic deletion of neuroligins in glia cells that are co-cultured with human neurons. Our results show that early postnatal deletion of neuroligins from astrocytes in vivo has no detectable effect on cortical or hippocampal synapses and does not alter the cytoarchitecture of astrocytes when evaluated in young adult mice. Moreover, deletion of astrocytic neuroligins in co-cultures of human neurons produced no detectable consequences for the formation and function of synapses. Thus, astrocytic neuroligins are unlikely to fundamentally shape synapse formation or astrocyte morphogenesis but likely perform other important roles that remain to be discovered.

## INTRODUCTION

Astrocytes perform vital roles across the lifespan of a synapse, from synapse formation to synaptic transmission to synapse elimination (reviewed in Farhy-Tselnicker and Allen, 2018; Eroglu and Barres 2010; Shan et al., 2021; Verkhratsky and Nedergaard, 2018; Lyon and Allen, 2022; Nagai et al., 2021). Indeed, at least a subset of synapses physically interacts with astrocytes to form ‘tripartite synapses’. The abundance for such astrocyte-synapse contacts varies among synapse types from <25% of synapses to nearly all synapses (Ventura and Harris, 1999; Araque et al., 1999; Kuwajima et al., 2013; Ostroff et al., 2014; Arizono et al., 2022; Salmon et al., 2023). Some synapses are always associated with astrocytic processes (e.g., parallel-fiber synapses in the cerebellum; Castejón, 1990; Baude et al., 1994), whereas others are never associated with astrocytes (e.g., calyx synapses in the brainstem; Meinrenken et al., 2003), and most are in between these two extremes.

The emergence of novel tools to genetically access and interrogate astrocytes has paved the way for a new effort dedicated to understanding the functional significance and the molecular mechanisms of the synaptic roles of astrocytes (Srinivasan et al., 2016; Yu et al., 2020, Gleichman et al., 2023, Shigetomi et al., 2013). Among others, RNA sequencing (RNAseq) studies have revealed astrocytic expression of cell-adhesion molecules that were traditionally viewed as specifically synaptic in nature (Saunders et al., 2018; Zeisel et al., 2018; Schaum et al., 2018; Zhang et al., 2014; Chai et al., 2017; Srinivasan et al., 2016; Clarke et al., 2018). The astrocytic expression of these synaptic cell-adhesion molecules gave rise to a parsimonious and attractive hypothesis that accounts for how astrocytes may interact with synapses at the molecular level (Stogsdill et al. 2017; Ackerman et al., 2021; Imrie et al., 2024; Chung et al., 2024; Saint-Martin and Goda, 2022; Tan and Eroglu, 2021). This hypothesis posits that astrocytes are integrated into tripartite synapses via synaptic cell-adhesion molecules, thereby enabling astrocytes to regulate synapse formation, maturation, function, or elimination.

Prominent among synaptic cell-adhesion molecules that are expressed by astrocytes are neuroligins, a family of postsynaptic adhesion molecules that are encoded in vertebrates by four genes (*Nlgn1-4*) (see Figure 1 below). The four neuroligin genes produce highly homologous proteins with an identical domain structure and a high degree of sequence similarity (Ichtchenko et al., 1995 and 1996; Bolliger et al. 2008). Neuroligins were discovered as the postsynaptic ligands for presynaptic neurexins but also bind to presynaptic LAR-type receptor tyrosine phosphatases and postsynaptic MDGA proteins (Liu et al., 2022; Liu et al., 2023; Connor et al., 2019; Qin et al., 2020; Südhof 2008; Bemben et al., 2015; Yoshida et al., 2021). Despite their similarity, however, different neuroligins display distinct localizations in brain and perform different non-redundant functions (Zhang et al., 2015; Chanda et al., 2018). Specifically, neuronal *Nlgn1* and *Nlgn2* function exclusively at excitatory and inhibitory synapses, respectively, whereas *Nlgn3* and Nlgn4 may act at both excitatory and inhibitory synapses (Ali et al., 2020; Qin et al., 2020; Connor et al., 2019; Krueger et al., 2012).

**Figure 1:**
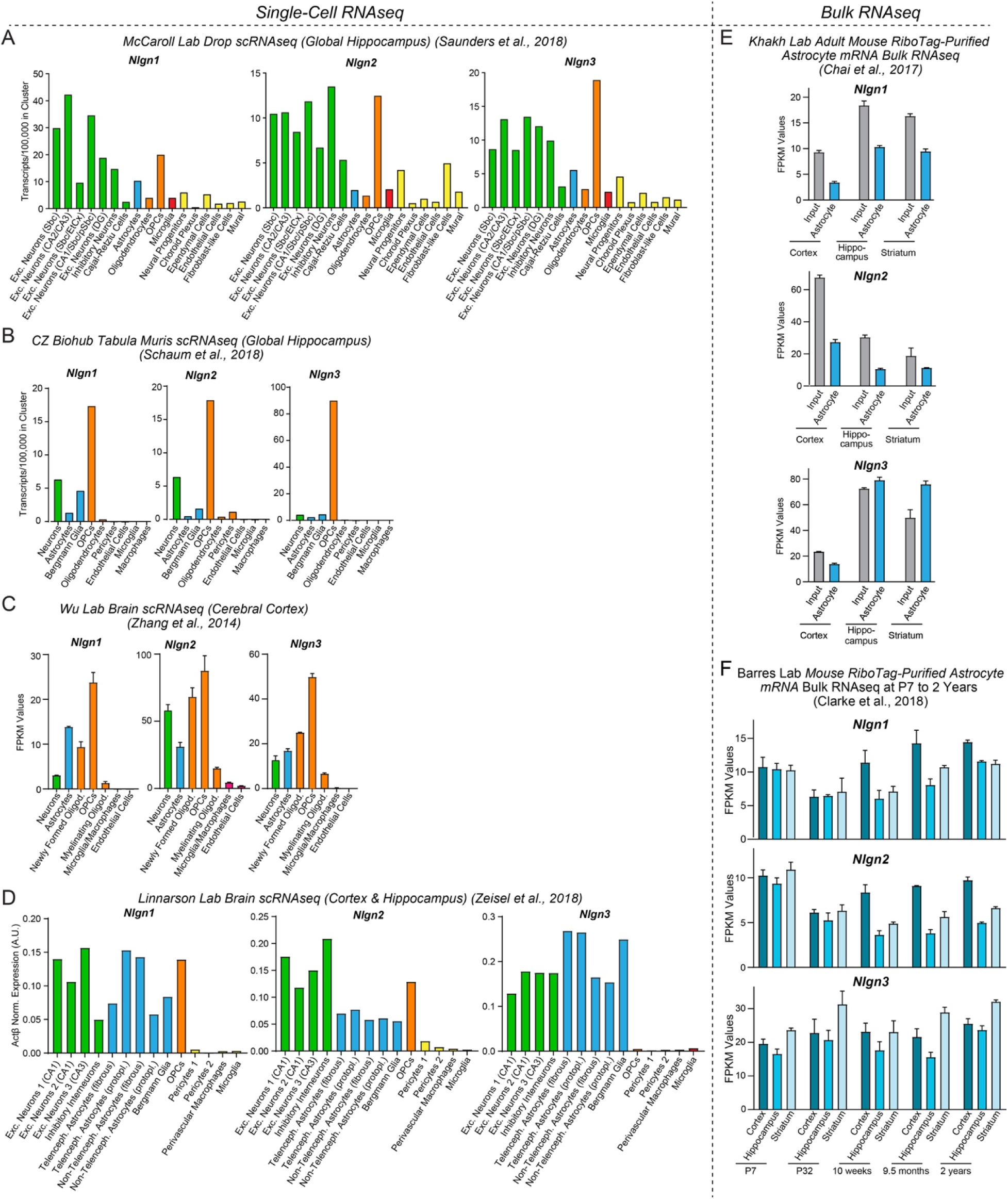
Neuroligin genes (*Nlgn1*-*3*) are abundantly expressed by neurons, astrocytes, and OPCs in brain as determined by analyses of publicly available RNAseq datasets. (**A-D**) Analysis of *Nlgn1*, *Nlgn2*, and *Nlgn3* mRNA levels in neurons (green), astrocytes (blue), oligodendrocyte lineage cells (orange), microglia (red), and other cell types in the brain (yellow) using the single-cell RNAseq dataset published from the McCaroll lab (Saunders and Macosko et al., 2018, www.dropviz.org) (**A**), Chan Zuckerberg Initiative (Schaum et al., 2018) (**B**), Wu lab (Zhang et al., 2014) (**C**), and Linnarson lab (Zeisel et al., 2018, www.mousebrain.org) (**D**). Note that although relative expression levels vary greatly between datasets, all datasets support the conclusion that Nlgn1, Nlgn2, and Nlgn3 are broadly expressed in astrocytes and OPCs. (**E**) Analysis of *Nlgn1*, *Nlgn2*, and *Nlgn3* mRNA levels in astrocytes in three different brain regions (cortex, hippocampus, and striatum) using the bulk RNAseq datasets published by the Khakh lab (Chai et al., 2017; Srinivasan et al., 2016, www.astrocyternaseq.org) that examined RiboTag-purified mRNAs. (**F**) Analysis of *Nlgn1*, *Nlgn2*, and *Nlgn3* mRNA levels in astrocytes as a function of age in mice using the bulk RNAseq datasets published by the Barres lab (Clarke et al., 2018; www.brainrnaseq.org). Astrocyte mRNA was purified from three different brain regions (cortex, hippocampus, and striatum) by RiboTag pulldowns in *Aldh1l1-eGFP-L10a* mice. Note that *Nlgn4* is not measured in the RNAseq experiments shown, probably because its expression levels are low and because its mRNA is very GC rich. For an analysis based on the pioneering original Barres lab data that used a less deep sequencing approach, see Figure S1.

In a tripartite synapse, astrocytic neuroligins would be ideally positioned to control synapse numbers or properties, for example by binding to presynaptic neurexins or postsynaptic MDGAs presented on the surface of synaptic membranes. Thereby, astrocytic neuroligins could regulate astrocyte-neuron interactions. Indeed, a landmark study concluded that astrocytic expression of neuroligins controls synapse formation in mice and that in particular one specific neuroligin isoform, *Nlgn2*, plays a central role in enabling excitatory synapse formation in the visual cortex (Stogsdill *et al*. 2017). Moreover, this paper described that *Nlgn2* and other neuroligins regulate the cytoarchitecture of astrocytes since a loss-of-function of *Nlgn2* and other neuroligins greatly reduced astrocyte branching in mixed cortical cultures of glia and neurons and in the visual cortex in vivo.

However, the impact of astrocytic *Nlgn2* deletions on excitatory synapse number reported by Stogsdill et al. (2017) was surprising because a preponderance of data indicates that *Nlgn2* protein (referred to as Nlgn2) is exclusively localized to, and functions in, inhibitory and not excitatory synapses. Specifically, multiple studies showed that in cultured neurons, Nlgn2 is only present in inhibitory synapses and that the deletion of *Nlgn2* causes a loss of inhibitory synapses and inhibitory synaptic responses but has no effect on excitatory synapse numbers or excitatory synaptic transmission (e.g., see Varoqueaux et al., 2004 and 2006; Graf et al., 2004; Chubykin et al., 2007; Poulopoulos et al., 2009; Chanda et al., 2019). These results thus showed that in culture, astrocytic or neuronal Nlgn2 has no essential role in excitatory synapse formation or function. Notably, in the same cultures the *Nlgn2* deletion did have a dramatic effect on inhibitory synapses. Moreover, classical experiments from the Barres lab revealed that astrocytes do contribute to synapse formation in culture preparations, suggesting that astrocytes are important for synapse formation even in culture preparations despite the limitations of such preparations (Pfrieger and Barres, 1997; Ullian et al., 2001).

It might be argued that the synaptogenic contribution of astrocytic Nlgn2 to excitatory synapse formation could have become dispensable in culture by an unknown mechanism, as opposed to a ‘real brain’ in which Nlgn2 might be required for excitatory synapse formation. Contrary to this argument, however, several *in vivo* studies of multiple brain regions confirm the conclusion from culture experiments that Nlgn2 is only present and only functions in inhibitory, but not in excitatory, synapses. Specifically, Patrizi et al. (2008) showed that in the cerebellar cortex, Nlgn2 protein exclusively localizes to inhibitory synapses even though excitatory synapses are vastly more abundant. No quantifications of the Nlgn2 localization were offered in this paper, presumably because no Nlgn2 signal was detectable in the numerous parallel-fiber and climbing-fiber synapses of the cerebellar cortex, whereas Nlgn2 was observed in 100% of inhibitory synapses. Blundell et al. (2009) quantified the density of excitatory and inhibitory synapses in *Nlgn2* constitutive KO mice in the adult hippocampal CA1 region using both confocal light microscopy and electron microscopy. In the mature brain, they found a decrease in inhibitory synapses (∼40%) but no change in excitatory synapses after constitutive germline deletions. Since the excitatory synapses in the CA1 region turn over every 2-3 weeks (Attardo et al., 2015; Pfeiffer et al., 2018), astrocytic Nlgn2 is clearly not essential in the hippocampus for excitatory synapse formation. These results also agree with the data of Panzanelli et al. (2017) who documented in the hippocampal CA1 region that Nlgn2 exclusively localizes to inhibitory synapses. Again, no quantifications were offered but the pictures are unequivocal in that staining for Nlgn2 overlaps 100% with that of two different markers for inhibitory synapse, gephyrin and GABA receptors. Furthermore, in likely the most definitive study to date, Kerti-Szigeti and Nusser (2016) performed a quantitative immuno-EM investigation of the GABA receptor composition of inhibitory hippocampal synapses. In the context of these studies, they labeled hippocampal sections with Nlgn2 antibodies. Although they also did not quantify excitatory synapses directly, their images and quantifications show that Nlgn2 completely co-localizes with GABA receptors at the EM level. These collective findings are mirrored by the experiments of Takacs et al. (2013) who show by immuno-EM that Nlgn2 is present in GABAergic and cholinergic synapses in the cortex, hippocampus, and several other brain regions. These authors also did not comment on excitatory synapses but inspection of their electron micrographic images indicates that Nlgn2 is absent from excitatory synapses. Finally, Poulopoulos et al. (2009) show that in the adult brainstem of constitutive Nlgn2 KO mice, inhibitory synaptic responses are severely impaired but excitatory synaptic responses are unchanged. Poulopoulos et al. (2009) also demonstrated that the constitutive *Nlgn2* KO suppresses hippocampal inhibitory synaptic responses but the authors again did not measure excitatory synaptic responses in the hippocampus.

Viewed together, the results from these prior studies are difficult to reconcile with the notion that astrocytic Nlgn2 is selectively essential for excitatory synapse formation as reported by Stogsdill et al. (2017). However, the many earlier studies cited here examined the constitutive *Nlgn2* deletion in which *Nlgn2* is deleted from neurons in addition to astrocytes, which may for an unknown reason cause a phenotype that is different from the astrocyte-only deletion of Stogsdill et al. (2017). It is thus conceivable that Nlgn2 has an essential function in excitatory synapse formation that only becomes manifest when *Nlgn2* is deleted from astrocytes but not neurons. Moreover, nearly all earlier studies did not examine the cortex but primarily studied the hippocampus, cerebellum, and brainstem, suggesting that Nlgn2 could perform distinct functions in the cortex and hippocampus. Finally, it is possible that differences in the timepoints of the deletions (i.e., constitutive deletions in the germline vs. in utero deletions) or in the developmental stages at which experiments were performed (young adult vs. adolescent mice) may have influenced the results.

Given the importance of understanding the potential role of astrocytic neuroligins in synapse formation and the differences between the results of Stogsdill et al. (2017) and of previous studies on Nlgn2, we have here examined the fundamental functions of astrocytic neuroligins. Using a rigorous genetic approach, we show that early selective postnatal deletions of *Nlgn1-3* in astrocytes do not significantly alter excitatory or inhibitory synapse numbers or functions (*Nlgn4* was not targeted owing to its low expression levels), at least when assessed in young adult mice. In addition, the astrocytic deletions of *Nlgn1-3* had no effect on the cytoarchitecture of astrocytes. Finally, deletion of all neuroligins (*Nlgn1-4*) in mouse glia that are then co-cultured with human neurons (which depend on the glia for synapse formation) did not detectably impair the ability of the mouse glia to promote neuronal synapse formation. Thus, astrocytic neuroligins in general and astrocytic Nlgn2 in particular likely do not perform a fundamental and essential function in promoting synapse formation or in shaping the astrocyte cytoarchitecture, suggesting that other astrocytic proteins are key drivers of synapse formation and that astrocytic neuroligins perform other important non-synaptic roles.

## RESULTS

### Astrocytes abundantly express neuroligins

For a gene to be involved in a particular physiological process, it must be expressed in the right place at the right time. To assess the expression of neuroligins in astrocytes in comparison with other cell types in brain, we examined the mRNA levels of *Nlgn1-3* in various brain cells by analysis of published RNA sequencing (RNAseq) datasets (Figure 1). *Nlgn4* mRNAs were not detected in these datasets, possibly because of the low expression levels of *Nlgn4* mRNAs or their high GC content. We quantified *Nlgn1*, *Nlgn2*, and *Nlgn3* mRNA transcripts in astrocytes and other cell types in the mouse hippocampus and cortex in the single-cell RNAseq datasets from the McCarroll Lab (Saunders and Macosko et al., 2018, www.dropviz.org), Chan Zuckerberg Initiative (Schaum et al. 2018), Wu lab (Zhang et al. 2014), and Linnarson lab (Zeisel et al., 2018, www.mousebrain.org) (Figure 1A-D). Interestingly, these quantifications showed that all three neuroligins were abundantly expressed in astrocytes and oligodendrocyte precursor cells (OPCs) in addition to neurons (Figure 1A-D). The relative expression levels of *Nlgn1*, *Nlgn2*, and *Nlgn3* in various cell types differed between datasets, probably because different cell isolation methods, RNAseq procedures and data analysis algorithms were used, but all datasets revealed similarly high neuroligin expression levels in neurons and astrocytes and generally even higher expression levels in OPCs. *Nlgn3* was consistently the most astrocyte-enriched neuroligin isoform (Figure 1A). The prominent expression of neuroligins in astrocytes was further supported by bulk RNAseq experiments performed by the Khakh lab (Chai et al., 2017; Srinivasan et al., 2016, www.astrocyternaseq.org) using astrocytic mRNAs purified via the RiboTag approach (Heiman et al., 2008) from the cortex, hippocampus, or striatum (Figure 1E). Again, all three isoforms were detected across these brain regions, with *Nlgn3* identified as the most enriched isoform compared to the input.

In mice, the major period for developmental synaptogenesis occurs over the first three weeks of life (Semple et al., 2013), although synapses are continuously formed and eliminated throughout life. Measurements of the expression of neuroligin proteins in brain as a function of postnatal development showed that neuroligin expression parallels the process of synaptogenesis (Irie et al., 1997). Moreover, comparative analysis of RNAseq data from the Barres lab (Clarke et al., 2018, www.brainrnaseq.org) indicates that during the entire lifespan of a mouse, astrocytes in the cortex, hippocampus, and striatum continuously express *Nlgn1-3* at similar levels (Figure 1F), consistent with a role of astrocytic neuroligins in the regulation of neural circuits. Thus, neuroligins are expressed in the right place and time for a role as astrocytic drivers of synapse formation.

### Conditional deletion of *Nlgn1-3* in astrocytes

To assess the fundamental functions of astrocytic neuroligins in synaptogenesis and astrocyte development, we generated mice that enable the inducible conditional deletion of astrocytic *Nlgn1*, *Nlgn2*, and *Nlgn3*. We targeted the conditional deletion specifically to astrocytes by crossing Aldh1l1-CreER^T2^ BAC transgenic mice that exhibit astrocyte-specific expression of tamoxifen-activatable Cre-recombinase (Winchenbach et al., 2016; Yu et al., 2020) with *Nlgn1*-*3* conditional KO (cKO) mice (Zhang et al., 2015) (Figure 2A). We then bred homozygous female triple *Nlgn1-3* cKO mice with male triple cKO mice carrying the Aldh1l1-CreER^T2^ allele to generate littermate male and female Cre-positive test mice and Cre-negative control mice. Mice were injected with tamoxifen intraperitoneally at postnatal days 10 and 11 (P10 and P11) or subcutaneously at postnatal day 1 (P1) and were analyzed at P35 – P48 (Figure 2B, 2C). The P10-11 tamoxifen injection paradigm was chosen based on a similar injection window used by Stogsdill et al. (2017) in *Nlgn2* cKO mice harboring GLAST-CreERT2 alleles, which resulted in a striking reduction in excitatory synaptic transmission in V1 layer 5 pyramidal neurons. On the other hand, the P1 injection was chosen to target astrocytes earlier in development and to better recapitulate the early postnatal targeting of astrocytes that Stogsdill et al. (2017) achieved using electroporation of P0/P1 mouse pups with plasmids to express shRNA (i.e. targeting individual *Nlgn1-3*) or Cre recombinase (in mice carrying floxed-Nlgn2 alleles). The mice carrying the astrocyte-specific deletion of neuroligins will subsequently be referred to as astro-Nlgn123 cKO mice.

**Figure 2:**
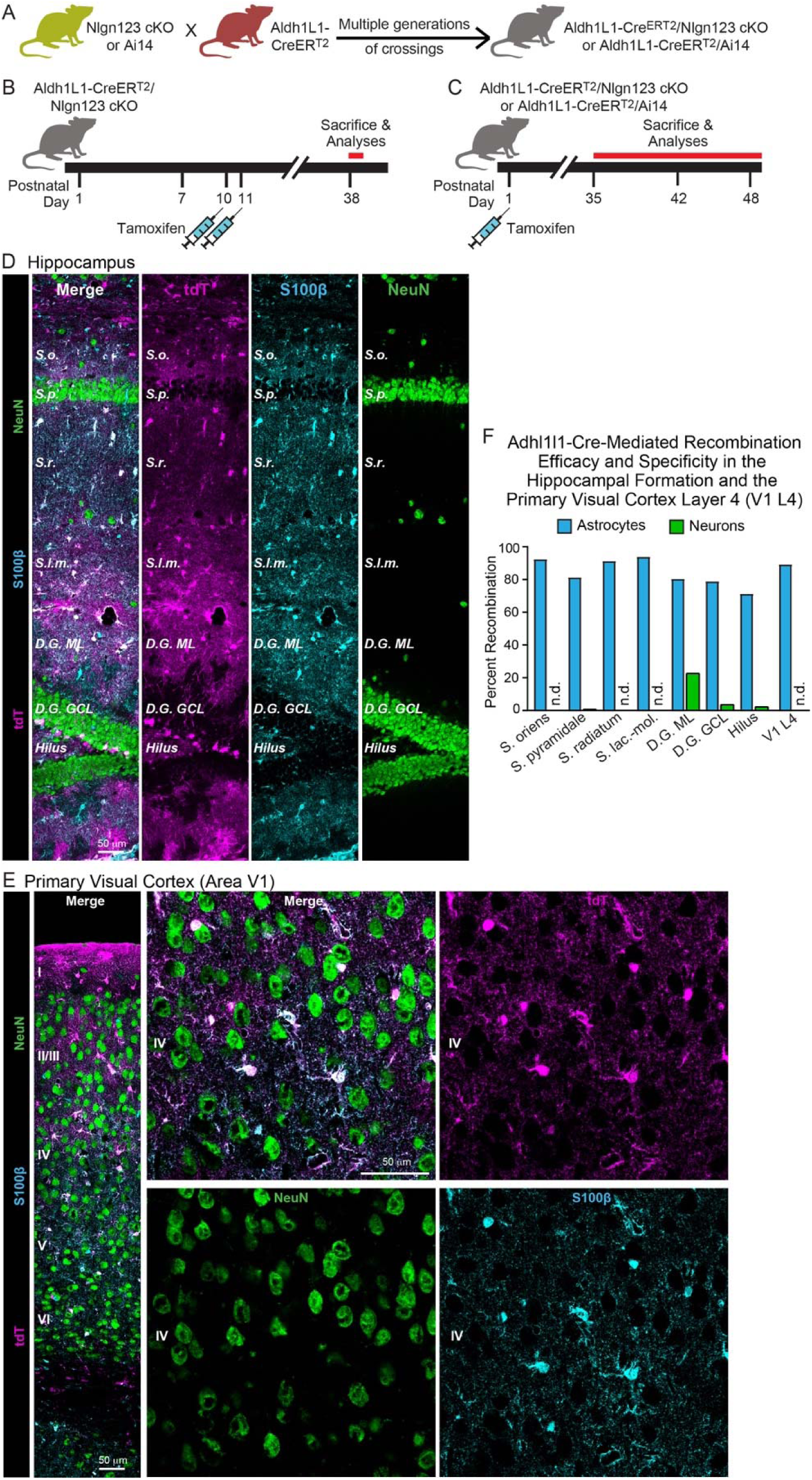
*Nlgn1*-*3* are efficiently and selectively deleted in astrocytes by crossing triple *Nlgn1-3* conditional KO mice with *Adh1l1*-CreER^T2^ driver mice and inducing Cre-activity with tamoxifen early during postnatal development. (**A**) Breeding strategy. Triple conditional KO mice carrying floxed *Nlgn1*, *Nlgn2*, and *Nlgn3* alleles or mice with a Cre-sensitive tdTomato (tdT) reporter allele (Ai14) were crossed with pan-astrocyte, tamoxifen-inducible Aldh1l1-CreER^T2^ BAC transgenic mice. *Nlgn1-3* cKO mice were crossed for multiple generations until homozygosity was reached (females: Nlgn1^f/f^ 2^f/f^ 3^f/f^, males: Nlgn1^f/f^ 2^f/f^ 3^f/y^). (**B** & **C**) Two different tamoxifen-induced Cre-activation schedules were used to delete *Nlgn1-3* in astrocytes. Aldh1l1-CreER^T2^ mice and controls (littermate Nlgn1-3 cKO mice lacking the Aldh1l1-CreER^T2^ allele) were injected with tamoxifen at P10 and P11 (**B**) (Trotter et al., 2021) or at P1 (**C**). Mice were sacrificed at least four (B) or five weeks post Cre induction to ensure complete deletion of neuroligins and decay of any astrocyte-specific neuroligin proteins. (**D** & **E**) To confirm the specificity and efficiency of the deletion of target genes in astrocytes using the Aldh1l1-CreER^T2^ BAC transgenic mouse line via tamoxifen injection at P1, Cre-recombination was visualized in the hippocampus (**D**) and primary visual cortex (**E**) via expression of tdT in reporter mice (magenta). Sections were additionally labeled for NeuN to mark neurons (green) and S100β to mark astrocytes (blue). (**F**) The Algh1l1-CreER^T2^-induced deletion of floxed genes produced by P1 tamoxifen injections is effective and selective for astrocytes as quantified using expression of tdTomato in reporter mice. tdTomato expression was quantified in the CA1 region of the hippocampus (the *S. oriens, S. pyramidale, S. radiatum,* and *S. lacunosum-moleculare*), the dentate gyrus (molecular layer [ML], granule cell layer [GCL], and hilus), and layer IV of the primary visual cortex.

### Efficient deletion of *Nlgn1-3* following P1 subcutaneous tamoxifen injection

To confirm the efficacy of CreER^T2^ induction, we focused on the subcutaneous tamoxifen injections at P1 since this condition was used for the majority of our experiments as the most rigorous approach but was not validated in detail previously. We crossed Aldh1l1-CreER^T2^ BAC transgenic mice with Cre-dependent tdTomato (tdT) reporter mice (Ai14), injected the resulting double-transgenic mice subcutaneously with tamoxifen at P1, and analyzed the mice at P35 by staining brain sections for tdTomato (Figure 2C). In the CA1 region of the hippocampus of these mice, we observed tdTomato-labeling in over 80% of astrocytes in the *S. oriens*, *S. pyramidale*, *S. radiatum*, or *S. lacunosum*-*moleculare*. In the cortex and hippocampus we found no neuronal recombination except for low levels in the dentate gyrus that is populated by adult-born neurons (Figure 2D & 2F). In layer IV (L4) of the primary visual cortex (V1), we also detected Cre-dependent tdTomato expression only in astrocytes without non-specific recombination in neurons. Again, approximately 90% of S100beta-expressing cells expressed tdTomato as an indicator of Cre activity (Figure 2E & 2F). Since some of the S100beta-expressing are OPCs, the true efficiency of Cre-mediated recombination likely exceeds 90%. Thus, the P1 tamoxifen injection protocol efficiently activates Cre recombinase in Aldh1l1-CreER^T2^ BAC transgenic mice.

We next determined the efficacy of the actual genetic recombination of *Nlgn1-3* alleles by employing a viral method that allows highly efficient capture of ribosome-bound mRNA (i.e. Ribotag) from infected astrocytes while minimizing unspecific capture of other mRNA’s, especially from neurons (Figure 3). This was accomplished by packaging in AAV2/5 an expression construct containing four elements: 1) the GfaABC1D promoter, which allows efficient expression of genetic cargo in astrocytes, 2) RiboTag (i.e., Rpl22-HA, which allows affinity capture of ribosome-bound mRNA with HA antibodies), 3) a membrane-targeted fluorescent reporter (i.e., Lck-mVenus), and 4) a 4×6T cassette of mIR-124 targeting sequences as was recently published (Gleichman et al., 2023). Inclusion of the 4×6T was essential to minimize off-target expression of RiboTag in neurons, which we had detected to occur in low but detectable levels in earlier versions of this construct lacking this cassette.

**Figure 3:**
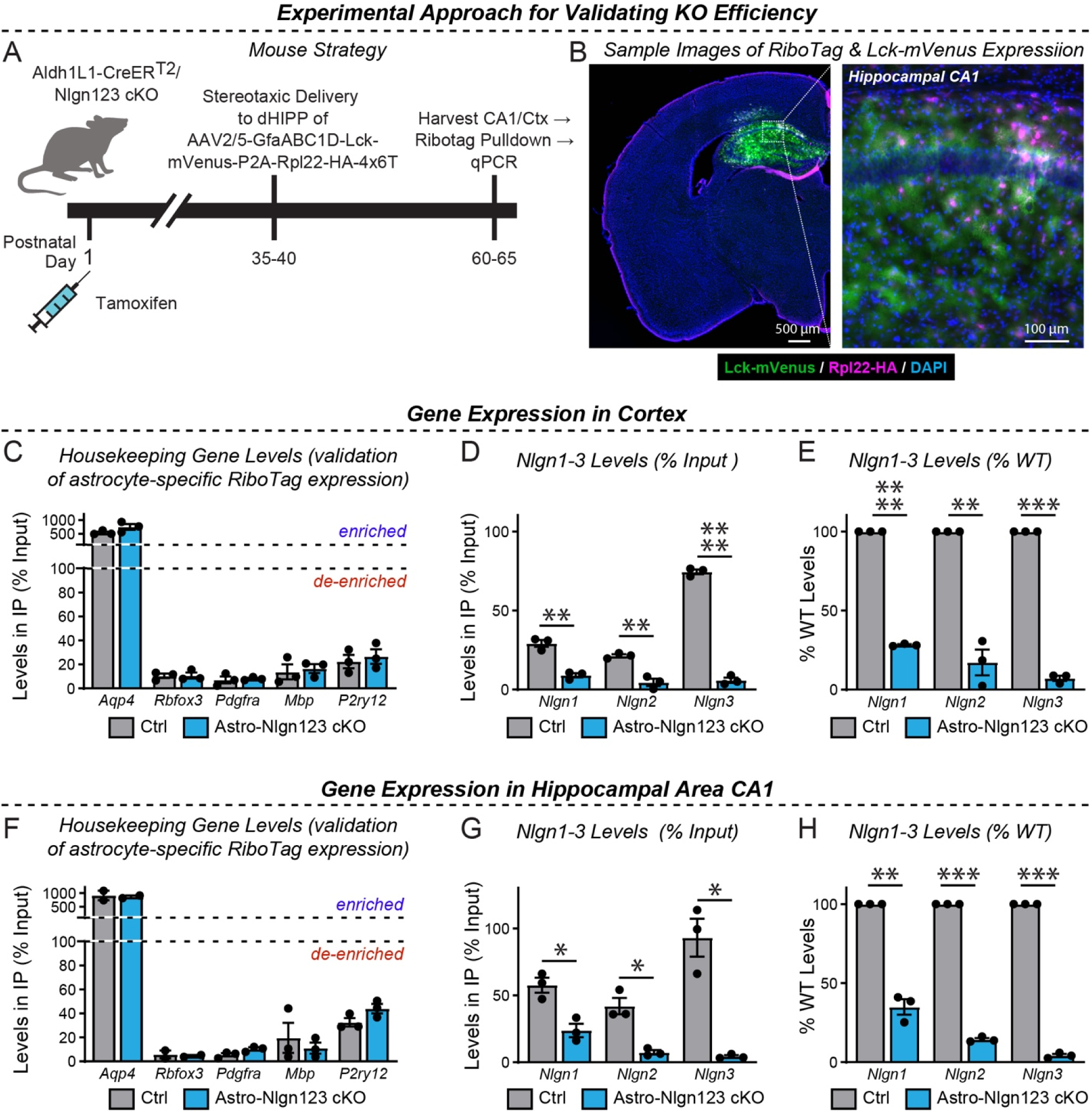
Quantification of the Nlgn1-3 conditional KO efficiency in astrocytes by qRT-PCR using RiboTag-mediated isolation of astrocytic mRNAs. (A) Diagram of the experimental strategy. AAVs (AAV2/5) encoding the RiboTag (Rpl22-HA) and Lck-mVenus (a membrane-targeted fluorescent reporter) driven by the astrocyte-specific GfaABC1D promoter and additionally including a 4×6T cassette of mIR-124 targeting sequences (Gleichman et al., 2023) were injected into the CA1 region of mice with a P0 induction of the astrocyte-specific Nlgn1-3 KO or control mice at P35-40 and analyzed 4 weeks later. (B) Representative coronal brain section (counterstained with DAPI) showing successful viral targeting of CA1 and adjacent regions indicated by the presence of Lck-mVenus (green) and Rpl22-HA (purple) astrocytes. (**C-E**) quantitative RT-PCR validation of the successful pull-down of highly-enriched astrocyte ribosome-bound mRNA from micro-dissected cortical tissue (i.e. Rpl22-HA). Enrichment of astrocyte marker, *Aqp4*, and de-enrichment of other cell marker genes is expressed as levels in pull-down (i.e. IP) relative to total mRNA (i.e. input). **(D)** Confirmation in astro-Nlgn123 cKO mice of reduced levels of *Nlgn1*, *Nlgn2*, and *Nlgn3* using assays that recognized floxed exons here expressed as levels in IP relative to input. **(E)** Same as D, except expressed as a % change relative to wild-type based on the <u>2-ΔΔCt method.</u> (**F** - **H**) Same as C-E, but for the hippocampal CA1 region. Data in summary graphs are means ± SEM with each data point representing individual animals. Unpaired two-tailed t-tests were used to test statistical significance of data in D and G, whereas a one sample t-test was used to test statistical significance of data in E and H (*, p < 0.05; **, p < 0.01; ***, p < 0.001; ****, p<0.0001).

We validated this approach using astro-*Nlgn123* cKO mice and their littermate controls, both of which were injected subcutaneously with tamoxifen on P1 (Figure 3A). At 5-6 weeks of age, mice were injected with equal amounts of AAV’s into the dorsal hippocampus. Approximately 3 weeks after surgery, acute slices were prepared in ice-cold cutting solution. After confirmation of the presence of Lck-mVenus(+)-astrocytes, the dorsal CA1 and nearby cortex were micro-dissected followed by lysis in the presence of cycloheximide. qPCR was performed in RiboTag pull-downs compared to input (i.e. total RNA) to assess the relative levels of marker genes for major cell types (e.g. astrocytes, neurons, OPC’s, oligodendrocytes, and microglia). In both the cortex and hippocampal area CA1, we found the expected enrichment of astrocytic *Aqp4* (∼5-10 fold) and de-enrichment of other cell markers, including a ∼95% de-enrichment of the neuronal marker *Rbfox3* (Figure 3C-3D).

We then examined the expression of *Nlgn1-3* using qRT-PCR with primers that recognize the exons that are targeted in the astro-*Nlgn123* cKO mice in order to avoid measurements of mRNAs that do not undergo efficient nonsense-mediated decay. In the CA1 region, we observed robust loss of *Nlgn3* (∼96%), *Nlgn2* (∼86%), and *Nlgn1* (65%) gene expression. Similarly, in the cortex, we observed a equivalent robust loss of *Nlgn3* (93%), *Nlgn2* (83%), and *Nlgn1* (72%) expression. Given that our targeting of astrocytes based on Ai14 Cre-reporter mice was ∼90-99% (Figure 2), these reductions are striking and definitive. The existence of some residual neuroligin transcripts likely reflects the presence of a small population of astrocytes heterozygous or wildtype for *Nlgn2* and *Nlgn3* or contaminations in the RiboTag pulldowns. In contrast, *Nlgn1* appears more difficult to recombine and it is likely that while many astrocytes are *Nlgn2* and *Nlgn3* knockout cells, a significant number may be heterozygous or wildtype for *Nlgn1*.

### Deletion of astrocytic neuroligins has no major effect on synaptic proteins

As a first step towards testing whether astrocytic neuroligins are essential for synaptogenesis in the hippocampus or cortex, we screened for changes in synaptic protein levels in astro-*Nlgn123* cKO mice. We collected hippocampal and cortical lysates from the brains of astro-*Nlgn123* cKO mice and from their Cre-negative littermates after induction of Cre-recombinase at P10 and P11 (Figure 4A-D) or at P1 (Figure 4E-H). Lysates were collected at P35-38 and analyzed by quantitative immunoblotting using fluorescent secondary antibodies. Both the levels of synaptic proteins and of neuroligins were measured in comparison to loading controls.

**Figure 4:**
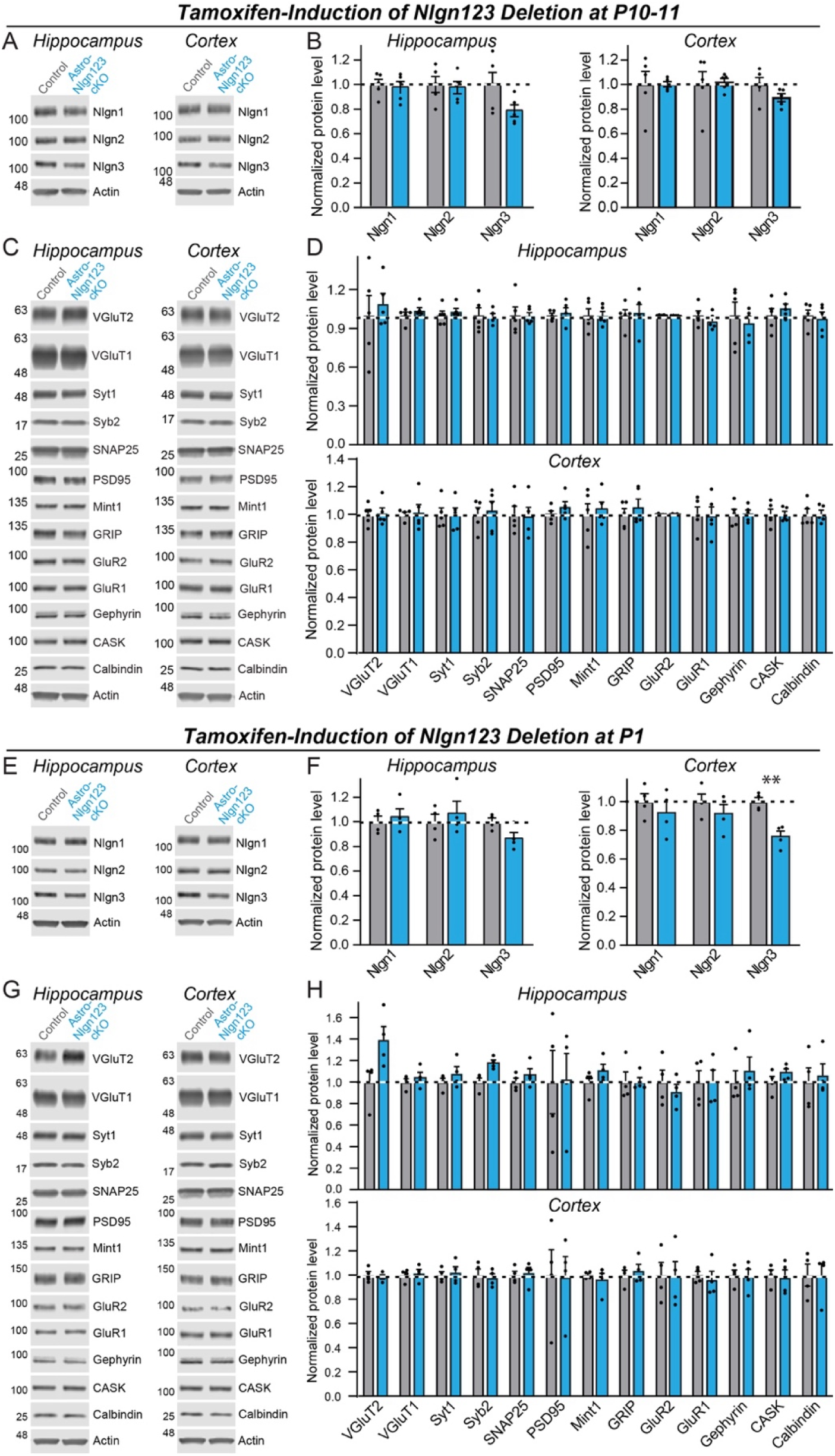
Conditional deletion of *Nlgn1*-*3* in astrocytes throughout the brain at early postnatal timepoints (P10/11 or P1) has only modest effects on overall neuroligin protein levels and does not significantly alter the synaptic proteome. (**A** & **B**) Representative immunoblots and quantifications of *Nlgn1*, *Nlgn2*, and *Nlgn3* protein levels from hippocampal (**A**) and cortical lysates (**B**) of astrocyte *Nlgn1-3* cKO and littermate control mice injected with tamoxifen at P10 and P11 and sacrificed at P38. Proteins were quantified on immunoblots using fluorescent secondary antibodies, with protein levels normalized to β-actin and then to control levels (n = 5, all male). (**C** & **D**) Representative immunoblots and (**D**) quantification for various synaptic protein levels from hippocampal and cortical lysates of astrocyte *Nlgn1-3* cKO and littermate control mice injected with tamoxifen at P10 and P11 and sacrificed at P38. Protein is quantified using fluorescent secondary antibodies, with protein levels normalized to β-actin and then to control levels (n = 5, all male). (**E-H**) Same as (**A-D**) except mice were injected with tamoxifen at P1 and sacrificed at P35 (n = 4, 2 male & 2 female). Numerical data are means ± SEM with statistical significance determined by unpaired two-tailed t-test (**, p<0.01).

We found that the levels of *Nlgn1*, *Nlgn2*, and *Nlgn3* proteins were not detectably decreased in either the cortex or the hippocampus after the P10-11 or the P1 induction of Cre-recombinase, except for a significant decline (∼25%) in *Nlgn3* protein levels in cortical lysates after the P1 induction (Figure 4A, B, E and F). Given that astrocytes account for only ∼20% of cells in brain, the absence of a significant decrease in neuroligin proteins in brain after the *Nlgn1-3* deletion in astrocytes is not surprising as one would expect maximally a ∼20% decrease in neuroligin levels if mRNA and protein levels perfectly correlated. Protein level measurements are inherently noisy, making detections of change of <20% difficult. The finding of a ∼25% decrease in *Nlgn3* levels in cortex after the P1 induction (Figure 4F) is consistent with this assessment since *Nlgn3* is the most abundantly expressed neuroligin isoform in astrocytes (Figure 1).

We next quantified the levels of selected synaptic proteins as a function of the astrocytic *Nlgn1-3* deletion (Figure 4C, D, G, and H). We analyzed 12 synaptic proteins as well as calbindin as a marker of subsets of inhibitory neurons. No significant changes in any protein analyzed were detected in the hippocampus or cortex. Thus, based on protein level measurements, no significant change in synapses was observed.

### Astrocytic neuroligins are dispensable for hippocampal synapse formation

Measurements of synaptic proteins in tissue lysates are a relatively insensitive approach to assessing synapse numbers. For a more direct measurement, we labeled cryosections from the brains of littermate astro-*Nlgn123* cKO and control mice at P35 after P1 induction of Cre-recombinase with antibodies to the excitatory presynaptic marker vGluT1, the excitatory postsynaptic marker Homer1 and the dendritic marker MAP2, and counterstained the sections with DAPI (Figure 5A). Because excitatory synapses in the hippocampus are too dense to be individually resolved by confocal imaging, we used the overall staining intensity as a proxy for synapse density. Low magnification (20x) imaging across the layers of the CA1 region of the hippocampus and the molecular layer of the dentate gyrus revealed no effect of the astro-*Nlgn123* cKO on vGluT1 or Homer1 staining intensity (Figure 5B). In order to increase the signal-to-noise ratio and improve our ability to detect small changes in staining intensity, we additionally performed high-magnification (60x) confocal imaging of the *S. pyramidale* and *S. radiatum* in the CA1 region (Figure 5A). Again, vGluT1 and Homer1 staining intensities were not altered by loss of astrocytic *Nlgn1*-3 at P1 (Figure 5B, C). Next, we stained hippocampal sections with antibodies to the excitatory presynaptic marker vGlut2 alongside Homer1 and MAP2 since vGluT2 is present only in a subset of excitatory synapses (Figure 5D). We also detected no changes in the vGluT2 staining intensity at either low or high magnifications (Figure 5E, F).

**Figure 5:**
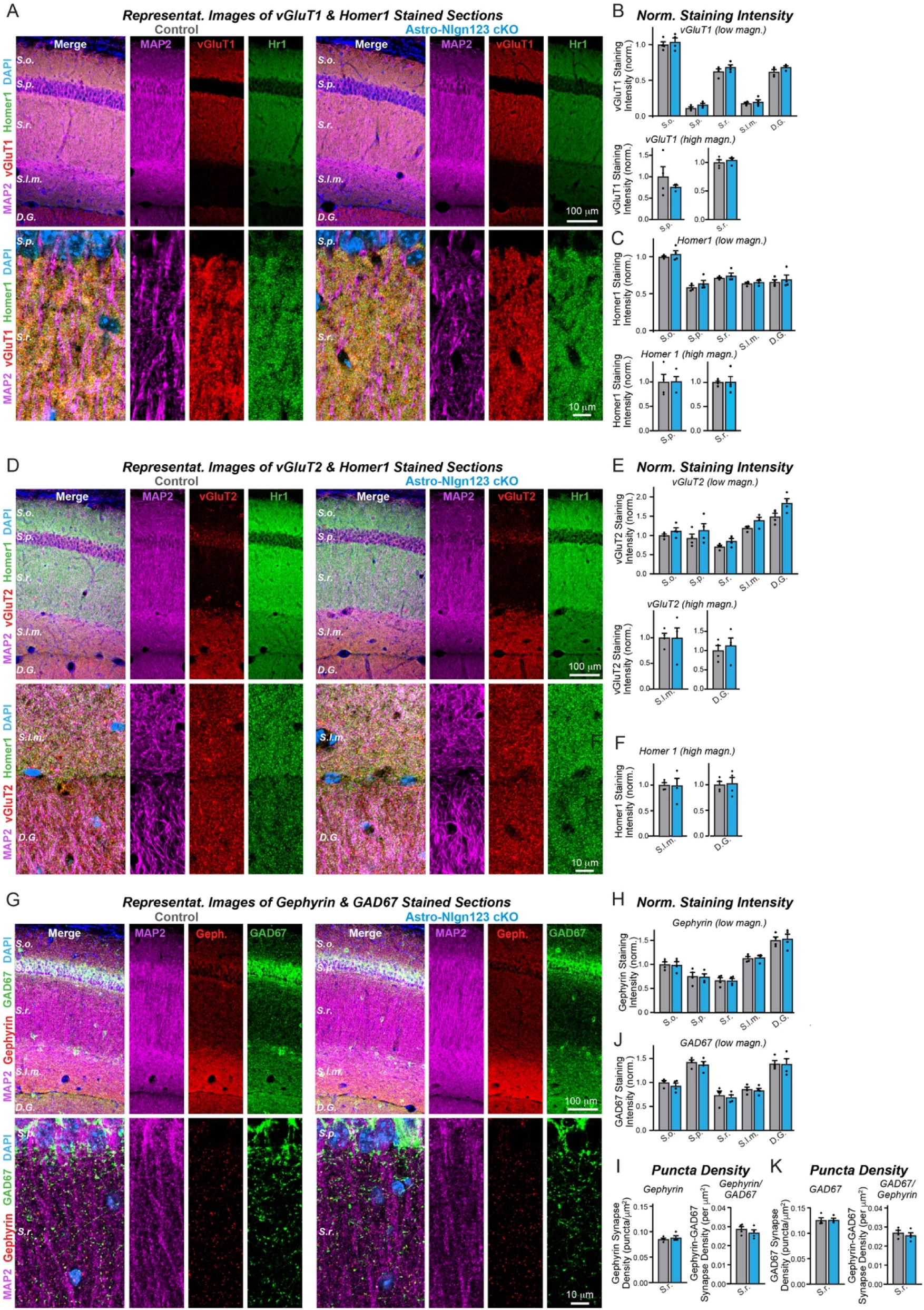
Conditional deletion of *Nlgn1*-*3* in astrocytes starting at P1 does not alter excitatory or inhibitory synapse numbers in the hippocampus as assessed by immunocytochemistry with antibodies to synaptic markers. (**A**) Representative images of CA1 and dentate gyrus hippocampal sections from astrocyte Nlgn1-3 cKO and littermate control mice, injected with tamoxifen at P1 and sacrificed at P35, stained for dendritic marker MAP2 (magenta), excitatory presynaptic marker vGluT1 (red), excitatory postsynaptic marker Homer1 (green), and DAPI (blue), taken at 20X (top) and 60X (bottom) magnification. (**B**) Quantification of total vGluT1 (top) and Homer1 (bottom) immunofluorescence signal for low magnification imaging (20X) across the layers of the hippocampus (*S. oriens, S. pyramidale, S. radiatum, S. lacunosum-moleculare,* dentate gyrus molecular layer), first internally normalized to MAP2 and then to average vGluT1 (top) or Homer1 (bottom) immunofluorescence level in *S. oriens* of control mice. (**C**) Quantification of total vGluT1 (top) and Homer1 (bottom) immunofluorescence signal for high magnification imaging (60X) in the CA1 *S. pyramidale* (left) and *S. radiatum* (right), first internally normalized to MAP2 and then to average vGluT1 (top) or Homer1 (bottom) immunofluorescence level in control mice. (**D**) Representative images of CA1 and dentate gyrus hippocampal sections from astrocyte Nlgn1-3 cKO and littermate control mice stained for dendritic marker MAP2 (magenta), excitatory presynaptic marker vGluT2 (red), excitatory postsynaptic marker Homer1 (green), and DAPI (blue), taken at 20X (top) and 60X (bottom) magnification. (**E**) Quantification of total vGluT2 immunofluorescence signal for low magnification imaging (20X) across the layers of the hippocampus, first internally normalized to MAP2 and then to average vGluT2 immunofluorescence level in *S. oriens* of control mice. (**F**) Quantification of total vGluT2 (top) and Homer1 (bottom) immunofluorescence signal for high magnification imaging (60X) in the CA1 *S. lacunosum-moleculare* (left) and dentate gyrus molecular layer (right), first internally normalized to MAP2 and then to average vGluT2 (top) or Homer1 (bottom) immunofluorescence level in control mice. (**G**) Representative images of CA1 and dentate gyrus hippocampal sections from astrocyte *Nlgn1-3* cKO and littermate control mice stained for dendritic marker MAP2 (magenta), inhibitory postsynaptic marker Gephyrin (red), inhibitory presynaptic marker GAD67 (green), and DAPI (blue), taken at 20X (top) and 60X (bottom) magnification. (**H**) Quantification of total Gephyrin (top) and GAD67 (bottom) immunofluorescence signal for low magnification imaging (20X) across the layers of the hippocampus, first internally normalized to MAP2 and then to average Gephyrin (top) or GAD67 (bottom) immunofluorescence level in *S. oriens* of control mice. (**I**) Quantification of puncta density for Gephyrin (top left), GAD67 (bottom left), Gephyrin having GAD67 (top right), and GAD67 having Gephyrin (bottom right) for high magnification imaging (60X) in the CA1 *S. radiatum*. Data are means ± SEM with statistical significance determined by unpaired two-tailed t-test (n=4, 2 male & 2 female).

To assess whether astrocytic neuroligins are required for inhibitory synapse formation, we labeled hippocampal sections with antibodies to the inhibitory presynaptic marker GAD67, the inhibitory postsynaptic marker gephyrin and the dendritic marker MAP2, and again counterstained the sections with DAPI (Figure 5G). We also observed no changes in gephyrin or GAD67 staining intensity after P1 deletion of astrocytic *Nlgn1-3* using low or high magnification imaging (Figure 5H, S1A). Given the lower density of inhibitory synapses in the hippocampus, individual inhibitory synapses could be resolved with high magnification imaging. This enabled us to quantify the density and size of the inhibitory GAD67 and Gephyrin synaptic puncta in the CA1 *S. radiatum*. Both were not changed in astro-*Nlgn123* cKO mice compared to control mice (Figure 5I, left panels and S1B). Finally, the number of synapses containing matched pre-(GAD67) and postsynaptic signals (gephyrin) was also quantified but exhibited no change in astro-*Nlgn123* cKO mice (Figure 5I and K, right panels). Thus, astrocytic *Nlgn1-3* are not essential for either excitatory or inhibitory synaptogenesis in the hippocampus of young adult mice.

### Astrocytic neuroligins are not required for basal synaptic function

The lack of a requirement of astrocytic neuroligins for synapse formation in the hippocampus, at least in young adult mice, agrees well with previous data demonstrating that constitutive neuroligin deletions do not decrease excitatory synapse numbers, but severely impair synaptic transmission (Varoqueaux et al., 2006; Chubykin et al., 2007; Poulopoulos et al., 2009; Patrizi et al., 2008; Blundell et al., 2009). To test whether loss of astrocytic *Nlgn1-3* causes a functional impairment of synapses, we monitored spontaneous excitatory and inhibitory synaptic transmission in CA1 region pyramidal neurons. We produced acute slices from littermate astro-*Nlgn123* cKO and control mice at P44-50 after P1 tamoxifen injections and performed patch-clamp recordings from CA1-region pyramidal neurons in the presence of tetrodotoxin (Figure 6). The amplitude, frequency, and kinetics of mEPSCs and mIPSCs were not changed by deletion of astrocytic *Nlgn1-3* (Figure 6). Furthermore, deletion of astrocytic neuroligins did not alter the membrane properties of CA1 pyramidal neurons (Figure S2). These data suggest that astrocytic neuroligins are not required for the development of functional synapses mediating basal synaptic transmission of CA1 pyramidal neurons.

**Figure 6:**
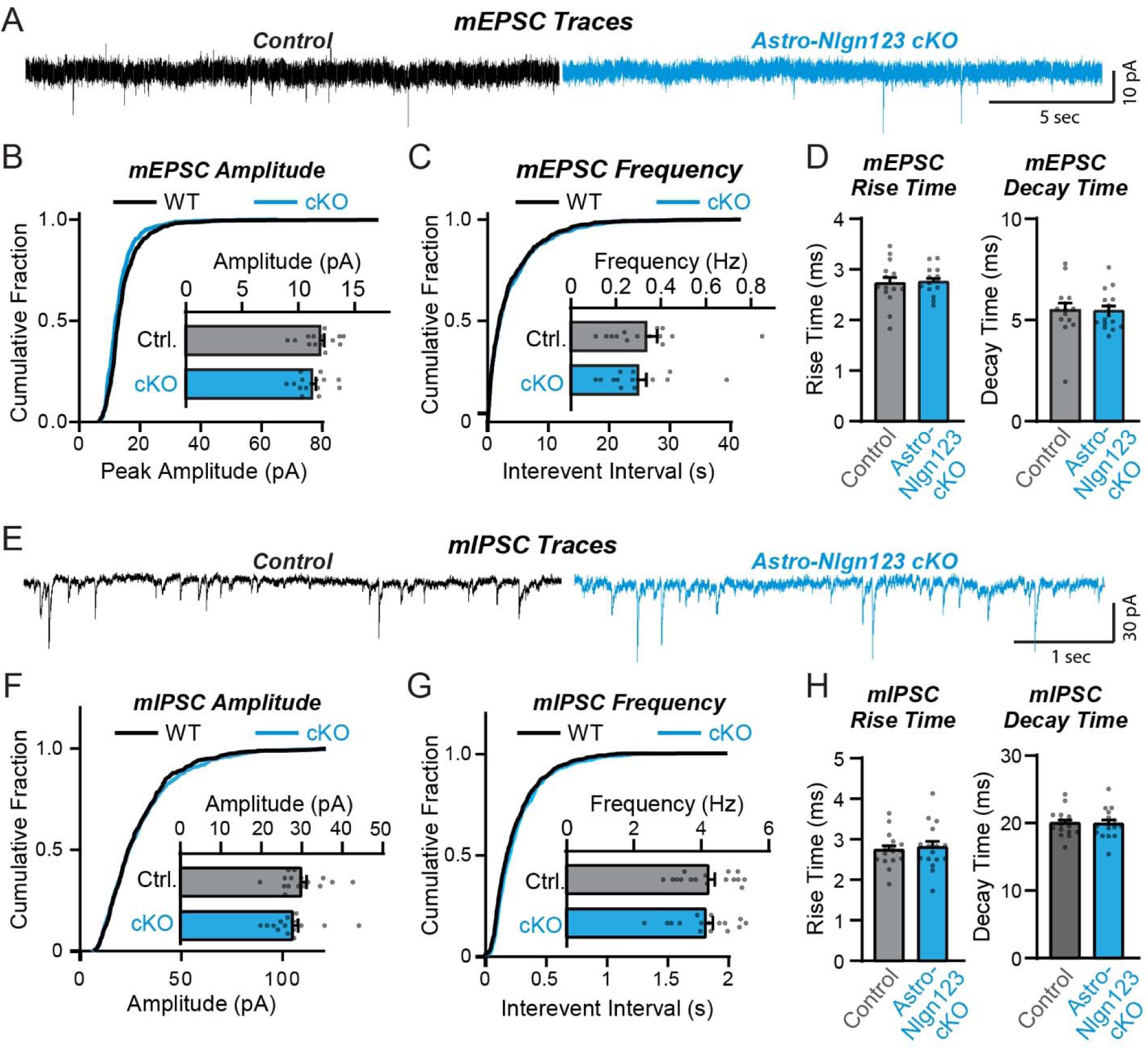
Conditional deletion of *Nlgn1*-*3* in astrocytes starting at P1 has no major effect on basal excitatory or inhibitory neurotransmission monitored in hippocampal CA1 pyramidal neurons. (A) Representative traces for miniature EPSCs (mEPSCs) from CA1 pyramidal neurons in acute slices from astrocyte *Nlgn1-3* cKO and littermate controls injected with tamoxifen at P1 and recorded at P44 - P50. (B) Cumulative distribution and summary graph of mEPSC amplitude and (**C**) frequency. (**D**) Summary graph of mEPSC rise (left) and decay (right) times (n = 14-15 cells / 3 mice per genotype). (**E-H**) Same as (**A-D**) except for miniature IPSCs (mIPSCs) (n = 16 cells / 3 mice per genotype). Data in summary graphs are means ± SEM with each data point representing individual cells. Unpaired two-tailed t-tests were used to test statistical significance of data in bar graphs, and Kolmogorov-Smirnov tests were used for cumulative curves (****, p<0.0001).

### Astrocytic neuroligins are not essential for synapse formation in the visual cortex

Since the finding of Stogsdill et al. (2017) of an essential function for astrocytic neuroligins in excitatory synapse formation was obtained in the visual cortex and it is possible that the function of astrocytic neuroligins differs between the hippocampus and the visual cortex, we explored the consequences of the genetic deletion of astrocytic *Nlgn1-3* on synapses in the visual cortex. We obtained cryosections of the visual cortex containing area V1 from astro-*Nlgn123* cKO and littermate control mice at P35 after the mice had been injected with tamoxifen at P1. The sections were co-stained with antibodies to Homer1, MAP2, and either vGluT1 or vGluT2 and counterstained with DAPI. We then imaged layer 4 (L4) of area V1 of the visual cortex, the same layer used by Stogsdill et al. (2017) (Figure 7A, C). As in the hippocampus, excitatory synapses in L4 of the primary visual cortex are too dense to be resolved individually, so we quantified the overall staining intensity as a proxy for synapse number. High magnification imaging revealed that loss of astrocytic neuroligins had no effect on the staining intensity of vGluT1 or vGluT2 but produced a small decrease (∼15%) in the staining intensity of Homer1 (Figure 7B, D).

**Figure 7:**
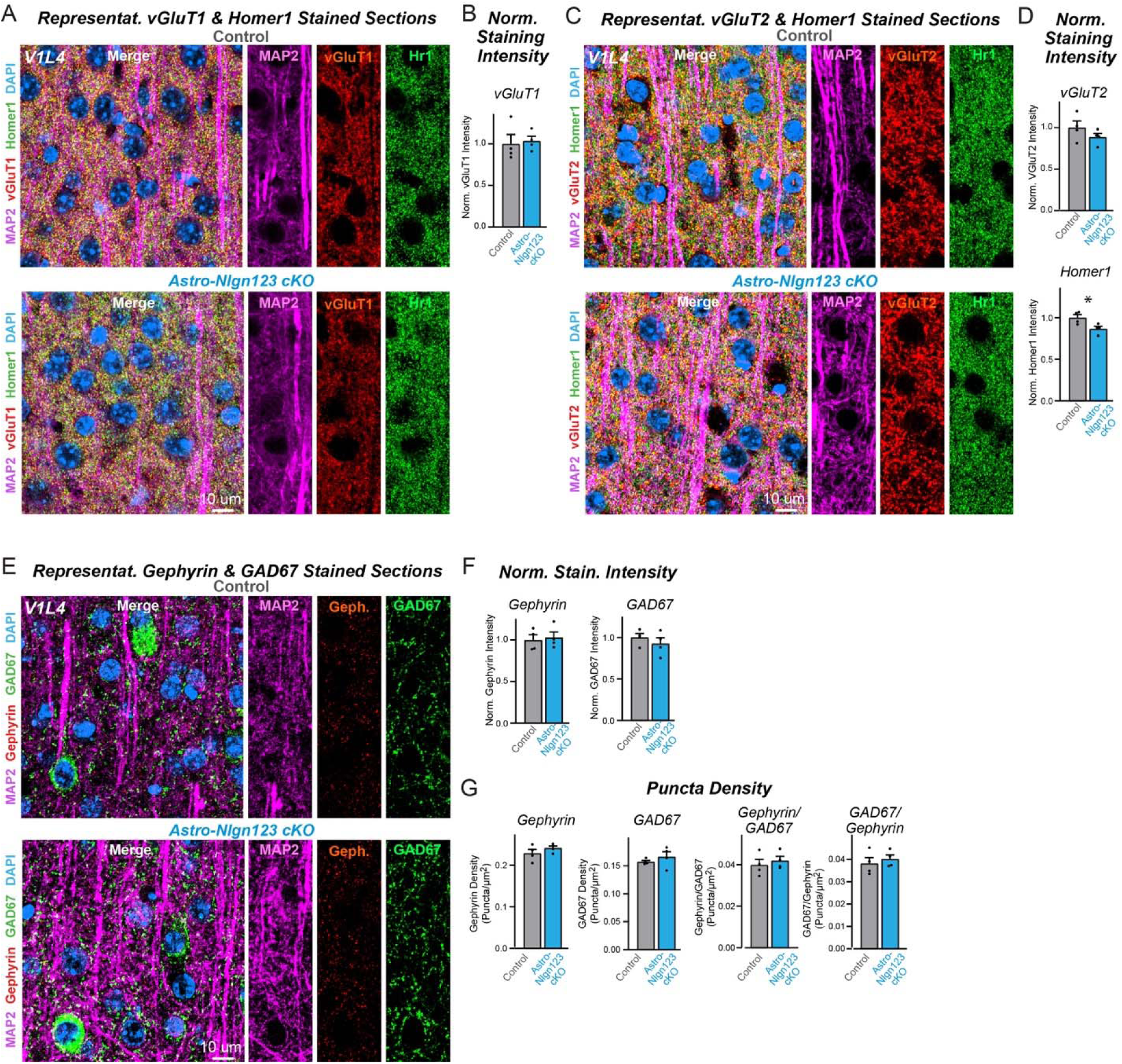
Conditional deletion of *Nlgn1*-*3* in astrocytes starting at P1 does not alter excitatory or inhibitory synapse numbers in layer IV of the primary visual cortex as assessed by immunocytochemistry with antibodies to synaptic markers. (**A**) Representative images of primary visual cortex (V1) layer IV (L4) astrocyte *Nlgn1-3* cKO and littermate control mice, injected with tamoxifen at P1 and sacrificed at P35, stained for dendritic marker MAP2 (magenta), excitatory presynaptic marker vGluT1 (red), excitatory postsynaptic marker Homer1 (green), and DAPI (blue) taken at 60X magnification. (**B**) Quantification of total vGluT1 (left) and Homer1 (right) immunofluorescence signal in V1L4 first internally normalized to MAP2 and then to average vGluT1 (left) or Homer1 (right) immunofluorescence level in control mice. (**C**) Representative images of V1L4 astrocyte *Nlgn1-3* cKO and littermate control mice stained for dendritic marker MAP2 (magenta), excitatory presynaptic marker vGluT2 (red), excitatory postsynaptic marker Homer1 (green), and DAPI (blue) taken at 60x magnification. (**D**) Quantification of total vGluT2 immunofluorescence signal in V1L4 first internally normalized to MAP2 and then to average vGluT2 immunofluorescence level in control mice. (**E**) Representative images of V1L4 astrocyte *Nlgn1-3* cKO and littermate control mice stained for dendritic marker MAP2 (magenta), inhibitory postsynaptic marker Gephyrin (red), inhibitory presynaptic marker GAD67 (green), and DAPI (blue) taken at 60X magnification. (**F**) Quantification of total Gephyrin (left) or GAD67 (right) immunofluorescence signal in V1L4 first internally normalized to MAP2 and then to average gephyrin (left) or GAD67 (right) immunofluorescence level in control mice. (**G**) Quantification of Gephyrin (left) or GAD67 (right) puncta density. (**H**) Quantification of Gephyrin (left) or GAD67 (right) puncta size in V1L4. (**I**) Quantification of puncta density for Gephyrin having GAD67 (left) or GAD67 having Gephyrin (right) in V1L4. Data are means ± SEM with statistical significance determined by unpaired two-tailed t-test (*, p<0.05) (n=4, 2 male & 2 female).

To assess the effect of deleting astrocytic neuroligins on inhibitory synapses in L4, sections were co-stained with antibodies to GAD67, gephyrin, and MAP2 and again counterstained with DAPI (Figure 7E). The staining intensity of GAD67 and gephyrin were unaffected by the astrocytic neuroligin deletion (Figure 7F). Since inhibitory puncta can be resolved with confocal imaging, we also quantified the density of GAD67 and gephyrin puncta (Figure 7G). Deletion of astrocytic *Nlgn1-3* also caused no change in any of these measures. These data suggest that, at least in our hands, astrocytic neuroligins are not fundamentally required for synaptogenesis in layer 4 of the primary visual cortex when measured at P35.

### Astrocytic neuroligins are not essential for astrocyte morphogenesis

It is possible that astrocytic neuroligins could be involved in shaping the cytoarchitecture of astrocytes as proposed by Stogsdill et al. (2017) even if they are not contributing to synapse number in the young adult brain. Thus, we asked whether astrocytic neuroligins contribute to the morphogenesis of astrocytes and/or the maintenance of their complex cytoarchitecture. We first measured the levels of a series of glial proteins in lysates of the hippocampus and cortex of astro-*Nlgn123* cKO and control mice that had been injected with tamoxifen at P1 and were analyzed at P35 (Figure 8A). However, we failed to uncover major changes (Figure 8B). Next, we immunostained astrocytes in CA1 hippocampal sections for glial fibrillary acidic protein (GFAP) that is constitutively expressed in mouse hippocampal astrocytes but did not detect any alterations in GFAP expression in astro-*Nlgn123* cKO mice at P35 (Figure 8C, D).

**Figure 8:**
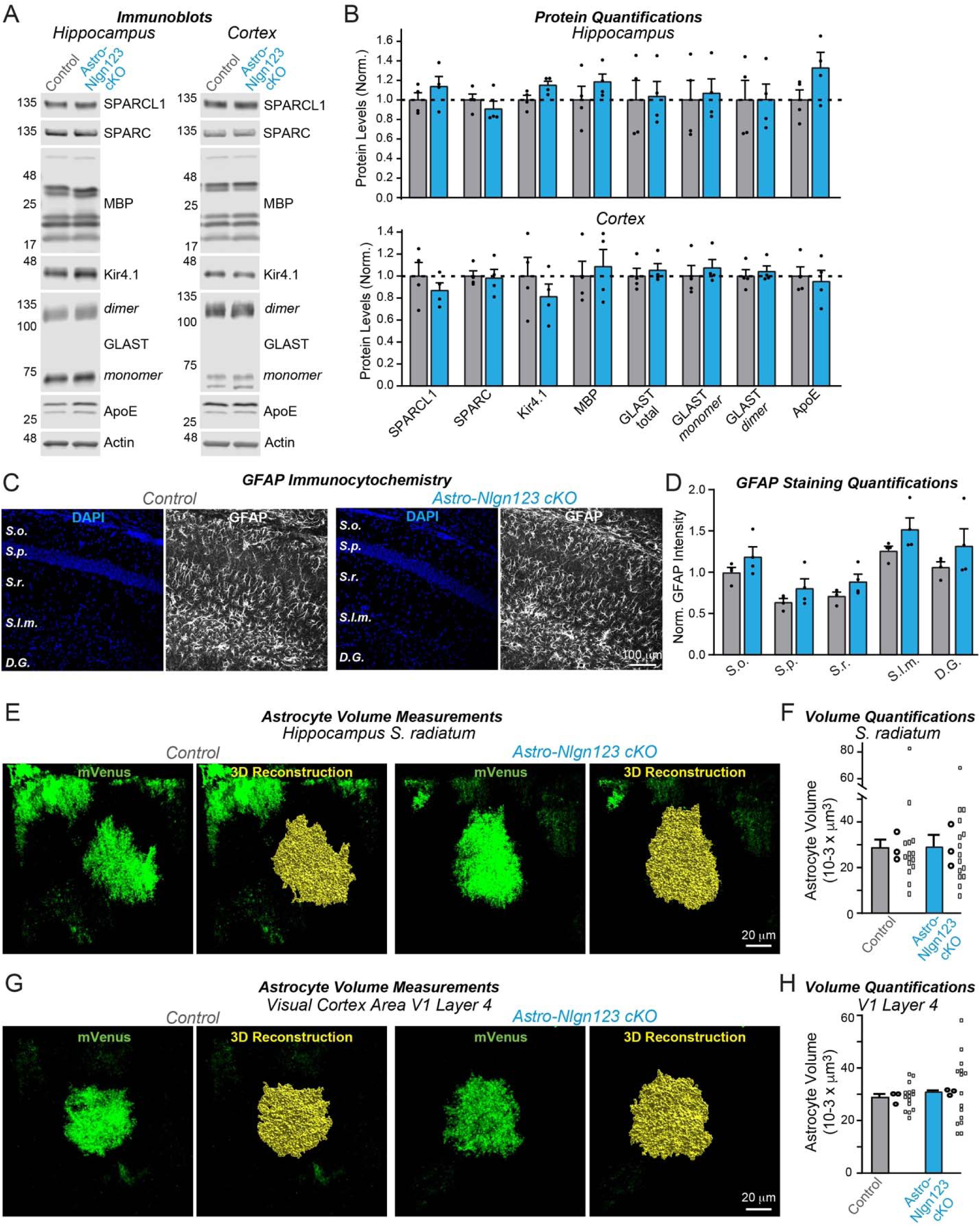
Conditional deletion of *Nlgn1*-*3* in astrocytes starting at P1 does not detectably alter the proteome or the cytoarchitecture of astrocytes in the hippocampus or layer IV of the primary visual cortex. (A) Representative immunoblots and (**B**) quantification for various glial protein levels from hippocampal and cortical lysates of astrocyte *Nlgn1-3* cKO and littermate control mice injected with tamoxifen at P1 and sacrificed at P35. Protein is quantified using fluorescent secondary antibodies, with protein levels normalized to β-actin and then to control levels (n = 4, 2 male & 2 female). (**C**) Representative images of CA1 and dentate gyrus hippocampal sections from astrocyte *Nlgn1-3* cKO and littermate control mice, injected with tamoxifen at P1 and sacrificed at P35, stained for astrocytic marker GFAP (white) and DAPI (blue), taken at 20X magnification. (**D**) Quantification of total GFAP immunofluorescence signal across the layers of the hippocampus (*S. oriens, S. pyramidale, S. radiatum, S. lacunosum-moleculare,* dentate gyrus molecular layer), normalized to average GFAP immunofluorescence level in *S. oriens* of control mice (n = 4, 2 male & 2 female). (**E**) To measure astrocyte volume, astrocyte *Nlgn1-3* cKO and littermate control mice were injected with tamoxifen at P1, underwent stereotactic injections of AAV expressing membrane-bound mVenus in astrocytes in the hippocampus at P14, and were subsequently sacrificed two weeks later at P35. Representative images of mVenus-expressing astrocytes in CA1 *S. radiatum* are shown with corresponding 3D volume reconstruction performed in Imaris. (**F**) Summary graph of CA1 *S. radiatum* astrocyte volumes shown averaged per animal with means ± SEM on the bar graph, as well as data points for individual astrocyte volumes. Statistical significance determined by unpaired two-tailed t-test of data averaged per animal (n=3, 1 male & 2 female). (**G & H**) Same as E & F, except for primary visual cortex layer IV astrocytes. Numerical data are means ± SEM. Dots in bar graphs represent independent biological replicates; in F and H, larger dots are independent biological replicates and smaller dots are pseudoreplicates since these are commonly reported in papers to boost statistical significance.

Finally, to directly test the claim that astrocytic neuroligins control astrocyte size (Stogsdill et al., 2017), we applied tamoxifen to astro-*Nlgn123* cKO and control mice at P1 and stereotactically injected AAVs expressing membrane-targeted mVenus under control of the GFAP promoter into their hippocampus or primary visual cortex at P21. We then imaged relatively thick sections (100 μm) from the hippocampus or primary visual cortex of these mice by confocal microscopy and reconstructed the entire volumes of astrocytes in the *S. radiatum* of the hippocampal CA1 region and in L4 of the primary visual cortex (Figure 8E, G). Quantifications of these volumes did not uncover any differences between astro-*Nlgn123* cKO and control mice, indicating that deletion of astrocytic *Nlgn1-3* did not alter astrocyte size or shape when analyzed at P35 (Figure 8F, H).

### Glial neuroligins are dispensable for synapse formation in co-cultured neurons

Human neurons are efficiently produced from human embryonic or induced stem cells, but only form reasonably mature synapses when co-cultured with glia cells that are usually produced from mouse embryos (Zhang et al. 2013; Huang et al., 2017). We thus asked whether neuroligins in mouse glia are contributing to the synaptogenic effect of glia co-cultured with human neurons (Figure 9A). We infected glia cultured from newborn mice with lentiviruses expressing inactive (ΔCre) or active Cre-recombinase (Cre), then added human neurons trans-differentiated from embryonic stem cells to the glia, and finally monitored synaptic connectivity using mEPSC measurements as a proxy for the combined effects of synapse numbers and synapse function (Figure 9A, B).

**Figure 9:**
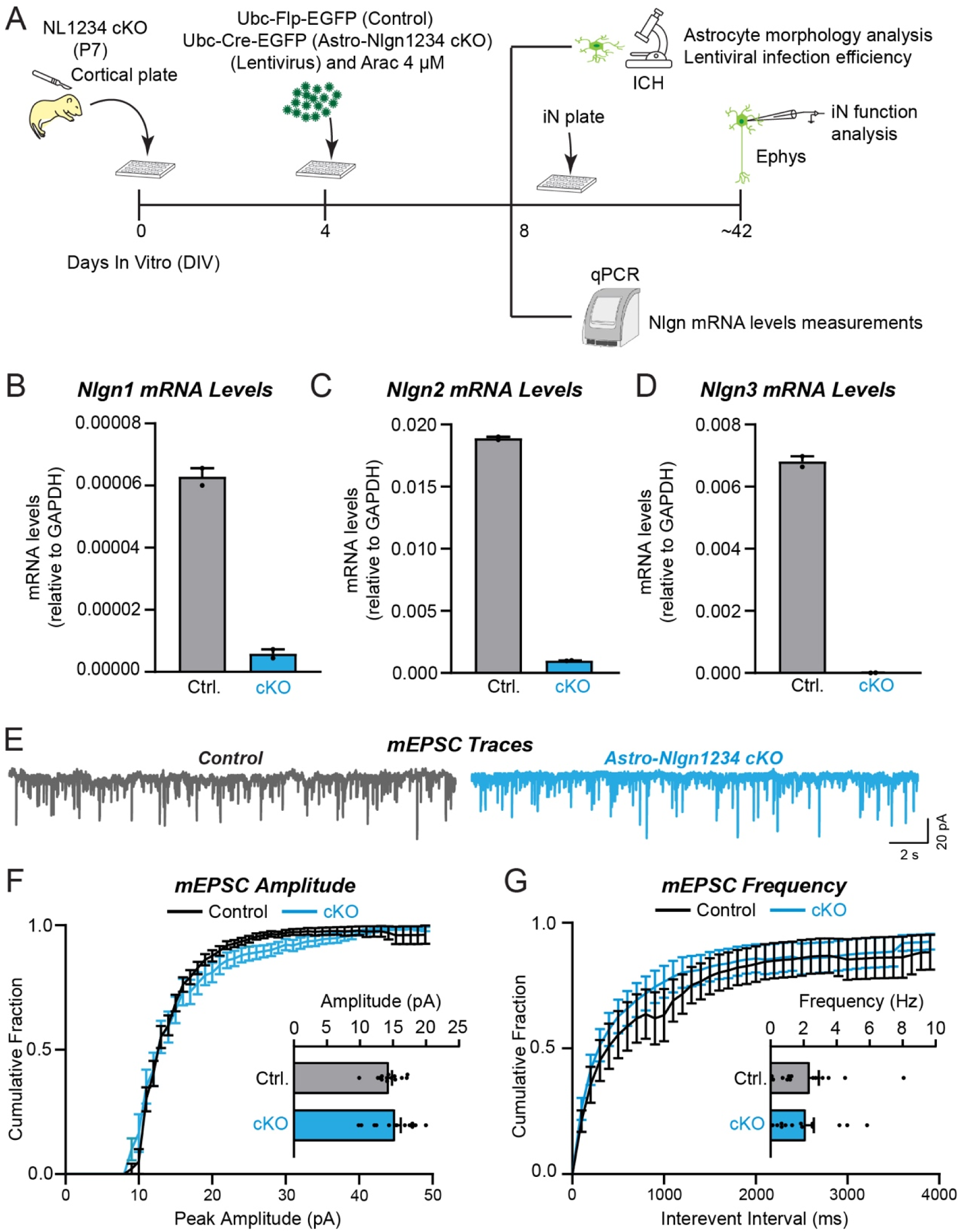
Deletion of *Nlgn1*-*4* in mouse glia co-cultured with wild-type human neurons does not significantly alter spontaneous synaptic events (mEPSCs) mediated by the co-cultured neurons. (**A**) Schematic of the experimental strategy (**B**-**D**) Measurements of the mouse Nlgn1, Nlgn2, and Nlgn3 mRNA levels in the mouse glia culture as a function of the expression of ΔCre (Ctrl., negative control) or Cre (cKO) reveals complete deletion of the expression of all neuroligins. (**E**-**G**) mEPSC measurements in human neurons co-cultured with mouse glia expressing (Ctrl.) or lacking neuroligins (cKO) fails to uncover any changes as a function of the neuroligin expression in the glia (E, sample traces; F, cumulative probability plot of the mEPSC amplitudes [inset, summary graph of the mean mEPSC amplitude]; G, cumulative probability plot of the mEPSC interevent intervals [inset, summary graph of the mean mEPSC frequency]. Numerical data are means ± SEM. Dots in bar graphs in F and G represent independent biological replicates.

qRT-PCR measurements confirmed that the deletion of all neuroligins in the glia was nearly complete (Figure 9B-D). Thus, the glia used in the test condition (Cre) express no detectable neuroligins. Quantifications of the frequency or amplitude of mEPSCs, however, demonstrated that the glial deletions of all neuroligins had no detectable effect on these parameters, suggesting that synaptic connectivity was unchanged (Figure 9E-G). These results indicate that the astrocytic neuroligins in dissociated mouse glia are not contributing to synapse formation when co-cultured with human neurons, even though previous experiments demonstrated that mouse glia greatly enhance synapse formation when co-cultured with human neurons (Zhang et al., 2013; Huang et al., 2017).

## DISCUSSION

Astrocytes are integral to shaping neural circuits. In the gray matter of the cerebral cortex (Halassa et al. 2007), hippocampus (CA1 *S. radiatum*) (Bushong et al. 2001), and cerebellar cortex (Spacek 1985), astrocytes (or their cousins, the Bergmann glia of the cerebellum) occupy non-overlapping territories, effectively “tiling” the neuropil. Within their territories, astrocytes not only contact all other cells in brain, but also associate with axons and dendrites and elaborate cellular processes close to synaptic junctions, often referred to as leaflets or perisynaptic astrocytic processes, thus forming tripartite synapses (Witcher et al., 2007; Derouiche et al., 2002; Imrie et al., 2024; Chung et al., 2024; Verkhratsky and Nedergaard, 2018; Saint-Martin and Goda, 2022; Oliveira and Araque, 2022; Lyon and Allen, 2022; Arizono and Nägerl, 2022). Astrocytes exhibit an exceptionally complex cytoarchitecture that includes thousands of fine processes infiltrating the neuropil where these processes likely perform multiple essential functions. At tripartite synapses, astrocyte functions include the removal of released glutamate from the perisynaptic area via glutamate transporters, thereby facilitating input specificity and preventing neurotoxicity (Rothstein et al., 1996; Tanaka et al, 1997; Yang et al., 2009; Bergles and Jahr, 1997; Bergles et al., 1997; Rose et al., 2018; Mahmoud et al., 2019). Astrocytes express numerous neurotransmitter receptors, enabling them to contribute to synaptic signaling and to respond to synaptic activity in a dynamic and activity-dependent manner (Tanaka et al., 1997; Tanaka et al., 2013; Yang et al., 2009; Allen and Eroglu, 2017). Moreover, astrocytic processes contain an array of ion channels that regulate their activity and contribute to ion homeostasis not only at synapses but also at other cellular locations, such as the nodes of Ranvier (Butt and Berry, 1994; Verkhratsky and Nedergaard, 2018; Black and Waxman, 1988) and blood vessels (Kuboter et al., 2019; Takano et al., 2006; Attwell et al., 2010). However, the molecular mechanisms by which astrocytes interact with synapses, blood vessels, nodes of Ranvier, and other brain components are incompletely understood.

Recent transcriptomic studies revealed that, in addition to neurotransmitter receptors and ion channels, astrocytes express a large array of cell-adhesion molecules that are known to function at synapses, suggesting that synaptic cell-adhesion molecules may mediate the integration of astrocytes into tripartite synapses and that astrocytes may regulate synapses via signals that are transmitted by such cell-adhesion molecules (Stogsdill et al. 2017; Ackerman et al., 2021; Imrie et al., 2024; Chung et al., 2024; Saint-Martin and Goda, 2022; Tan and Eroglu, 2021). Indeed, our analysis of multiple independent RNAseq datasets showed that astrocytes express particularly high levels of neuroligins, canonical postsynaptic cell-adhesion molecules that are known to be essential for synaptic function (Figure 1). The absolute and relative expression levels of different neuroligin isoforms in astrocytes varied among studies, but all studies concur in the conclusion that the mRNA levels of the three major neuroligins (*Nlgn1-3*) are overall similar in astrocytes and neurons (Figure 1; *Nlgn4* was not captured in the RNAseq studies owing to the high GC content of its mRNA and its lower abundance). Thus, neuroligins are clearly not neuron-specific, which is consistent with the possibility that neuroligins in astrocytes may embed astrocytes in tripartite synapses and enable astrocytes to contribute to the formation and performance of synapses.

Motivated by the observation of high levels of neuroligin expression in astrocytes and by the findings of Stogsdill et al. (2017) that suggest a major role for astrocytic neuroligins in synapse formation and have guided the understanding of the role of astrocytes at synapses, we here investigated the function of neuroligins expressed in astrocytes. We aimed to achieve two related goals: First, to test the hypothesis emerging from the Stogsdill et al. (2017) paper that astrocytic neuroligins as synaptic cell-adhesion molecules perform a critical role in promoting synapse formation. Second, to resolve a major contradiction arising from a comparison of the Stogsdill et al. (2017) findings with previous studies that argue against this hypothesis. Stogsdill et al. (2017) showed that astrocytic neuroligins in general are essential for excitatory synapse formation and for the normal morphology of astrocytes, and that *Nlgn2* in particular is specifically required for excitatory synapse formation. Inconsistent with this conclusion, however, earlier studies detected *Nlgn2* protein of any origin only in inhibitory but not in excitatory synapses (Varoqueaux et al., 2004; Graf et al., 2004; Poulopoulos et al., 2009; Patrizi et al., 2008; Blundell et al., 2009; Panzanelli et al., 2017; Takacs et al., 2013; Kerti-Szigeti and Nusser, 2016) and showed that deletion of *Nlgn2* in both neurons and glia caused a dysfunction of only inhibitory but not of excitatory synapses (Vareauqueaux et al., 2006; Chubykin et al., 2007; Poulopoulos et al., 2009; Blundell et al., 2009; Chanda et al., 2017). Since this contradiction remained unaddressed in studies following the Stogsdill et al. paper (2017), we felt that it was necessary to try to resolve it.

In order to investigate whether astrocytic neuroligins, as opposed to neuronal neuroligins, perform an essential function in synapse formation, we used a rigorous genetic approach. We generated a tamoxifen-inducible mouse model that enables conditional deletion of *Nlgn1-3* in all astrocytes (Figure 2A), which we confirmed allowed their efficient deletion (Figure 3). We then analyzed the effect of such deletion in depth in the hippocampal CA1 region and the visual cortex using two different time points of induction of the deletion of astrocytic neuroligins (P1 and P10-11; Figure 2). We made two principal observations.

First, deletion of *Nlgn1-3* from astrocytes had no detectable effect on the density of excitatory or inhibitory synapses in the hippocampus or cortex, or on the properties of spontaneous excitatory or inhibitory synaptic transmission in the hippocampus when measured in young adult mice. We established this conclusion by immunocytochemical analyses using a panel of antibodies to synaptic markers (Figure 5, 7), by electrophysiology (Figure 6), and by quantification of the synaptic proteome (Figure 4). Moreover, human neurons co-cultured with mouse glia do not depend on glial neuroligins for synapse formation, even though the human neurons require mouse glia for efficient synapse formation (Figure 9). Thus, astrocytic *Nlgn1-3* are not required for synapse formation and/or maintenance in mice at P35-42 or in co-cultures of human neurons with dissociated mouse glia.

Second, deletion of *Nlgn1-3* from astrocytes does not cause a measurable change in the cytoarchitecture of astrocytes (Figure 8). This was assessed by monitoring the astrocyte protein composition, GFAP staining in the hippocampus, and three-dimensional reconstruction of astrocytes in the hippocampus or cortex. Thus, *Nlgn1-3* are not fundamentally required for the morphogenesis or cell shape maintenance of astrocytes when analyzed in young adult mice.

Two questions arise at this point. Is it possible that the genetic deletion of *Nlgn1-3* in astrocytes for some reason was ineffective? To assess the efficiency of the tamoxifen-induced deletion of floxed genes, we quantified Cre-activity using the P1 induction protocol, which is the approach used for most experiments in this paper since it aims to permanently delete all neuroligins at a time early in development (Figure 2D-F). We observed Cre-mediated recombination in >80% of astrocytes but in <5% of neurons except for the dentate gyrus in which neurons are continuously replenished by adult neurogenesis. This finding confirms that the genetic approach efficiently induces Cre-recombinase activity, which in turn we previously demonstrated using the same conditional triple *Nlgn1-3* alleles employed here quantitatively deletes all three neuroligin genes (Chanda et al., 2017; Zhang et al., 2015). We further confirmed efficient recombination of *Nlgn1-3* using a viral RiboTag approach (Figure 3), which showed the highly efficacy knockout of *Nlgn2* and *Nlgn3*, and to a lesser extent *Nlgn1*. In addition, we demonstrated that the deletion of glial neuroligins in the co-culture experiments with human neurons was complete (Figure 9).

The second question is how can we explain the discrepancies between our results and those of Stogsdill et al. (2017)? It is possible that multiple differences in the approaches used may have contributed to these discrepancies.

First, we used a purely genetic approach, whereas a large amount of the data in the Stogsdill et al. (2017) paper were obtained with shRNAs. shRNA-mediated knockdowns are prone to off-target effects and additionally suffer from the limitation that their use interferes with the entire microRNA processing machinery of a cell. In addition, Stogsdill et al. (2017) used conditional *Nlgn2* KO mice (the same alleles that we originally generated and that were used in the present study) for analyses of synapse numbers, astrocyte morphology, and spontaneous synaptic transmission. To confirm *Nlgn2* shRNA findings, Stogsdill et al. (2017) employed postnatal astrocyte labeling by electroporation (PALE) to sparsely express Cre and a fluorescent reporter in astrocytes started on P2. They then confirmed a similar change in astrocyte morphology as was caused by shRNA, as well as ∼50% reductions in thalamocortical and intracortical excitatory synapses in the territories of V1 L4 astrocytes. They also used tamoxifen-induced deletion of astrocytic *Nlgn2* at P10-11 with the GLAST-CreER^T2^ mouse line, which led led to an impairment in the function of excitatory synapses in the layer V of the primary visual cortex (Stogsdill et al. 2017). Thus, although it is not possible with this approach to assess the efficiency of the *Nlgn2* deletion, at least some of the findings by Stogsdill et al. (2017) cannot be completely explained by RNAi-dependent off-target effects.

Second, Stogsdill et al. (2017) largely employed ‘sparse’ deletions of neuroligins in a subset of astrocytes whereas we used a global deletion of neuroligins in nearly all astrocytes, and that they mostly analyzed adolescent mice whereas we examined young adult mice. Sparse deletions, such as those used by Stogsdill et al. (2017), are powerful tools but their phenotypes are difficult to interpret because a manipulated cell with a deletion will be at a competitive disadvantage with surrounding wild-type cells. As a result, changes that are not related to the function of the deleted gene but are caused by this competitive disadvantage become manifest as phenotypes. It is thus possible, maybe even likely, that neuroligins perform a non-synaptic function in astrocytes, and that owing to indirect downstream effects induced by the competition with surrounding wild-type astrocytes, the shape of the mutant astrocytes and the formation of excitatory synapses may have been compromised even though astrocytic neuroligins are not important for the formation of synapses *per se*. Astrocytes tile the neuropil by engaging in a competition with neighboring astrocytes for individual territories through a mechanism similar to that of dendritic tiling (Freeman, 2010). Thus, we might expect that a sparse manipulation would produce a misleading phenotype over that of a global manipulation. An indirect effect produced by a competitive disadvantage induced with a sparse deletion has been recently reported in astrocytes for hepatocyte cell-adhesion molecule (hepaCAM). In this study, sparse deletion of hepaCAM led to a reduction in the volume of the astrocyte territory, while global deletion of hepaCAM had no effect (Baldwin et al. 2021), although there was an increase in astrocyte overlap in the global hepaCAM knockouts. Additionally, it has been reported that while the global deletion of *Nlgn1 in vivo* has no effect on synapse number in cortical layer 2/3 pyramidal neurons, sparse deletion via electroporation of shRNAs targeting *Nlgn1* does result in a reduction of spine density and synapse number (Kwon et al., 2012).

Third, we analyzed a different developmental timepoint than Stogsdill et al. (2017). Our analyses were performed in young adult mice (largely P35), whereas Stogsdill et al. (2017) examined synapses in adolescent mice (P21) when synaptic and astrocytic morphological refinements are advanced but not yet fully complete (Freeman 2010). Thus, it is possible that the phenotype observed by Stogsdill et al. (2017) is developmentally transient in nature and recovers in the time difference between their and our analysis. A transient effect of a neuroligin-like gene in astrocytes has been described in Drosophila. Here, RNAi knockdown of astrocytic Drosophila *Nlgn2* delayed motor circuit closure during development but did not result in robust, lasting behavioral phenotypes after the critical period had passed (Ackerman et al. 2021). It is important to note that, in agreement with our data indicating that astrocytic neuroligins do not control astrocyte morphogenesis, astrocyte-specific deletions of the Drosophila *Nlgn2* also had no effect on astrocyte volume or tiling (Ackerman et al. 2021). Furthermore, in a screen investigating astrocyte morphological diversity across the nervous system, neuroligins were not identified among the genes correlating with differences in astrocyte morphology (Endo et al. 2022).

In summary, it seems unlikely that astrocytic neuroligins have a fundamental and long-lasting function in synapse formation or in shaping the cytoarchitecture of astrocytes. However, it is probable that neuroligins perform other important roles in astrocytes that remain to be identified. Indeed, two recent studies on astrocytic *Nlgn3* have uncovered roles in social behavior (Dang et al., 2024; Qin et al., 2024). Continual advances in the tools available to access astrocytes, in concert with genetic models that allow temporally-defined manipulation of genes, will be key to discovering these roles that may provide new insights into astrocyte biology.

## EXPERIMENTAL PROCEDURES

### Key resources table

| Reagent type (species) or resource | Designation | Source or reference | Identifiers | Additional information |
| --- | --- | --- | --- | --- |
| Strain, strain background (Mus musculus) | Aldh111-Cre/ERT2 BAC transgenic mice | The Jackson Laboratory | RRID:IMSR_JAX:029655 |  |
| Strain, strain background (Mus musculus) | Nlgn123 cKO mice | Südhof lab |  |  |
| Cell line | HEK293T cells | ATCC | RRID: CVCL_0063 |  |
| Antibody | Anti-Nlgn1 | Synaptic Systems | 129111, RRID:AB_887747 | 1:2000 |
| Antibody | Anti-Nlgn2 | Synaptic Systems | 129203, RRID:AB_993014 | 1:1000 |
| Antibody | Anti-Nlgn3 | Synaptic Systems | 129311, RRID:AB_2151947 | 1:2000 |
| Antibody | Anti-β Actin | Sigma Aldrich | A1978, RRID:AB_476692 | 1:5000 |
| Antibody | Anti-VGluT2 | Millipore Sigma | AB2251, RRID:AB_1587626 | 1:1000 |
| Antibody | Anti-VGluT1 | TCS | YZ6089, RRID:AB_2861224 | 1:1000 |
| Antibody | Anti-Synaptotagmin 1 | TCS | V216 | 1:1000 |
| Antibody | Anti-Synaptobrevin 2 | TCS | P939 | 1:1000 |
| Antibody | Anti-SNAP25 | TCS | P913, RRID:AB_2861227 | 1:1000 |
| Antibody | Anti-PSD95 | Synaptic Systems | 124011, RRID:AB_10804286 | 1:1000 |
| Antibody | Anti-Mint1 | TCS | P730 | 1:1000 |
| Antibody | Anti-GRIP | BD Transduction Laboratories | 611319, RRID:AB_398845 | 1:1000 |
| Antibody | Anti-GluR2 | Neuromab | 75-002, RRID:AB_2232661 | 1:1000 |
| Antibody | Anti-GluR1 | Neuromab | 75-327, RRID:AB_2315840 | 1:1000 |
| Antibody | Anti-Gephyrin | Neuromab | 75-444, RRID:AB_2636852 | 1:1000 |
| Antibody | Anti-CASK | Neuromab | 75-000, RRID:AB_2068730 | 1:1000 |
| Antibody | Anti-Calbindin | Sigma | C9848, RRID:AB_476894 | 1:2000 |
| Antibody | Anti-Mouse SPARC-like1 | R&D Systems | BAF2836, RRID:AB_2195096 | 0.5 μg/ml |
| Antibody | Anti-SPARC | DSHB | mAB 236, RRID:AB_2617208 | 1 μg/ml |
| Antibody | Anti-Myelin Basic Protein | EnCOR Biotechnology | CPCA-MBP, RRID:AB_2572352 | 1:5000 |
| Antibody | Anti-Kir4.1 | Millipore Sigma | AB5818, RRID:AB_92053 | 1:1000 |
| Antibody | Anti-EEAT1 | Abcam | ab416, RRID:AB_304334 | 1:1000 |
| Antibody | Anti-MAP2 | EnCOR Biotechnology | CPCA-MAP2, RRID:AB_2138173 | 1:500 |
| Antibody | Anti-Gephyrin | Synaptic Systems | 147318, RRID:AB_2661777 | 1:200 |
| Antibody | Anti-GAD67 | Millipore | MAB5406, RRID:AB_2278725 | 1:1000 |
| Antibody | Anti-VGluT1 | Millipore | AB5905, RRID:AB_2301751 | 1:1000 |
| Antibody | Anti-MAP2 | Sigma Aldrich | M1406, RRID:AB_477171 | 1:1000 |
| Antibody | Anti-Homer1 | TCS | YZ6081 | 1:1000 |
| Antibody | Anti-NeuN | Millipore Sigma | MAB377, RRID:AB_2298772 | 1:1000 |
| Antibody | Anti-S100 | Abcam | ab868, RRID:AB_306716 | 1:1000 |
| Antibody | Anti-GFP | Invitrogen | A-11122, RRID:AB_221569 | 1:1000 |
| Antibody | Anti-HA.11 Epitope Tag-Alexa647 | BioLegend | 682404, RRID: AB_2566616 | 1:250 |
| Antibody | Alexa fluor 647, goat anti rabbit IgG | Invitrogen | A-21245, RRID:AB_141775 | 1:1000 |
| Antibody | Alexa fluor 546, goat anti rabbit IgG | Invitrogen | A-11035, RRID:AB_143051 | 1:1000 |
| Antibody | Alexa fluor 633, goat anti- mouse IgG | Invitrogen | A-21052, RRID:AB_2535719 | 1:1000 |
| Antibody | Alexa fluor 488, goat anti guinea pig IgG | Invitrogen | A11073, RRID:AB_2534117 | 1:1000 |
| Antibody | Alexa fluor 647, goat anti chicken IgG | Invitrogen | A21449, RRID:AB_1500594 | 1:1000 |
| Antibody | Alexa fluor 546, goat anti guinea pig IgG | Invitrogen | A-11074, RRID:AB_2534118 | 1:1000 |
| Antibody | Alexa fluor 488, goat anti mouse IgG | Invitrogen | A-11001, RRID:AB_2534069 | 1:1000 |
| Antibody | IRDye 800CW donkey anti chicken | Licor | 926-32218, RRID:AB_1850023 | 1:10000 |
| Antibody | IRDye 800CW donkey anti goat | Licor | 926-32214, RRID:AB_621846 | 1:10000 |
| Antibody | IRDye 680RD donkey anti guinea pig | Licor | 926-68077, RRID:AB_10956079 | 1:10000 |
| Antibody | IRDye 800CW donkey anti rabbit | Licor | 926-32213, RRID: AB_621848 | 1:10000 |
| Antibody | IRDye 680LT donkey anti mouse | Licor | 926-68022, RRID: AB_621848 | 1:10000 |
| Antibody | IRDye 800CW donkey anti mouse | Licor | 926-32212, RRID: AB_10715072 | 1:10000 |
| Chemical compound, drug | Picrotoxin | Tocris | 1128 |  |
| Chemical compound, drug | CNQX | Tocris | 1045 |  |
| Chemical compound, drug | D-AP5 | Tocris | 0106 |  |
| Chemical compound, drug | Tetrodotoxin | Cayman Chemical | 14964 |  |
| Chemical compound, drug | Tamoxifen | Sigma | T5648-1G |  |
| Chemical compound, drug | Corn Oil | Sigma | C8267 |  |
| Oligonucleotides | Actb PrimeTime qPCR Assay | IDT | Mm.PT.51.1402242 |  |
| Oligonucleotides | Aqp4 PrimeTime qPCR Assay | IDT | Mm.PT.58.9080805 |  |
| Oligonucleotides | Sox9 PrimeTime qPCR Assay | IDT | Mm.PT.58.42739087 |  |
| Oligonucleotides | Pdgfra PrimeTime qPCR Assay | IDT | Mm.PT.56a.5639577 |  |
| Oligonucleotides | MBP PrimeTime qPCR Assay | IDT | Mm.PT.58.28532164 |  |
| Oligonucleotides | P2ry12 PrimeTime qPCR Assay | IDT | Mm.PT.58.43542033 |  |
| Oligonucleotides | Rbfox3 PrimeTime qPCR Assay | IDT | Mm.PT.58.11398454 |  |
| Oligonucleotides | Nlgn1 PrimeTime qPCR Assay | IDT | Mm.PT.58.5704919 |  |
| Oligonucleotides | Nlgn2 PrimeTime qPCR Assay | IDT | Mm.pt.58.41142345 |  |
| Oligonucleotides | Nlgn3 PrimeTime qPCR Assay | IDT | Mm.PT.58.31138258 |  |
| Software, algorithm | Clampfit | Molecular Devices | N/A |  |
| Software, algorithm | pClamp | Molecular Devices | RRID:SCR_011323 |  |
| Software, algorithm | Prism | Graphpad software inc | RRID: SCR_002798 |  |
| Software, algorithm | Image Studio Lite | Licor | RRID: SCR_014211 |  |
| Software, algorithm | Imaris | Oxford Instruments | RRID:SCR_007370 |  |
| Software, algorithm | NIS-Elements Basic Research | Nikon | RRID:SCR_002776 |  |

#### Mice

Aldh1l1-Cre/ER^T2^ BAC transgenic mice were purchased from The Jackson Laboratory (Strain #:029655, RRID:IMSR_JAX:029655) and were bred with *Nlgn1*-*3* cKO mice, generated as previously described (Zhang et al., 2015), until homozygosity was reached for the floxed *Nlgn1-3* alleles. Mice were maintained on a mixed CD1 and C57BL/6 background. Litters resulting from crossing female *Nlgn1-3* cKO mice with male triple cKO mice carrying the Aldh1l1-CreER^T2^ were injected with tamoxifen at early postnatal timepoints to generate astro-*Nlgn123* cKO mice and littermate controls for experiments. After weaning between postnatal days 20 and 24, mice were housed in groups on a 12-hour light-dark cycle with open access to food and water. All mouse handling and procedures were conducted as approved by the Stanford University Administrative Panel on Laboratory Animal Care.

#### Tamoxifen Injections

To prepare tamoxifen for injection, 1g of tamoxifen (Sigma, Cat # T5648-1G) was mixed with 10 ml of 200 proof ethyl alcohol (Gold Shield) at room temperature for 15 minutes while protected from light with foil. This mixture was then combined with 90 ml of corn oil (Sigma, Cat# C8267) and agitated at 37°C for 1-2 hours until fully dissolved, while continuing to protect from light. Aliquots of 1 ml were stored at −20°C. On the day of injections, tamoxifen aliquots (10mg/ml) were thawed to room temperature. Tamoxifen solution was injected with an insulin syringe intraperitoneally/subcutaneously with a series of two 30 µl injections at P10 and P11 or one 20 µl injection subcutaneously in the neck at P1.

#### Immunoblotting

Mice were anesthetized with isoflurane and then decapitated. Hippocampus and cortex were collected in RIPA buffer (50mM Tris-HCl, 150 mM NaCl, 1% Triton X-100, 0.1% SDS, 1 mM EDTA) with cOmplete, EDTA-free protease inhibitor cocktail (Roche, Cat# 11873580001) on ice. Tissue samples were homogenized with a dounce tissue grinder on ice, rotated at 4°C for 30 min, and finally centrifuged at 14,000 rpm at 4°C for 20 min. Supernatant was collected from the samples and protein content was quantified using a BCA assay (Thermo, Cat# 23225). Protein lysates were stored at −80°C until use.

Quantified protein lysates were added to Laemmli sample buffer with DTT. Samples were boiled at 95°C for 5 min, except in the case of immunoblotting for multi-pass transmembrane proteins, for which samples were not heated. Samples were run on 4-20% Criterion TGX precast gels (Bio-Rad, Cat# 5671094 & 5671095) to separate proteins by molecular weight. Proteins were transferred from gels to 0.2 µm nitrocellulose membranes (Bio-Rad, Cat# 1620112) using the Trans-blot turbo system (Bio-Rad) for 7 min at 25V. For detection of proteins, membranes were first blocked in 5% non-fat milk (Carnation) in TBST for one hour at room temperature, followed by incubation with primary antibodies diluted in 5% non-fat milk (Carnation) in TBST overnight at 4°C Membranes were washed with TBST three times for 10 min per wash prior to incubation with LI-COR secondary antibodies diluted in 5% non-fat milk (Carnation) in TBST at a concentration of 1:10,000. Membranes were subsequently imaged with the Odyssey CLx imaging system (LI-COR) with quantification carried out in Image Studio Lite 5.2. Protein quantifications were normalized to beta-actin and to protein levels of controls, as described in applicable figure legends.

#### Immunohistochemistry

Mice were anesthetized with isoflurane and then pericardially perfused with phosphate buffered saline (PBS) for 1 min followed by ice cold 4% paraformaldehyde (PFA) for 7 min at a rate of 1 µl/min. PBS and PFA were filtered through a 0.2 µm filter prior to perfusion. Brains were extracted and post-fixed in 4% PFA overnight at 4°C. PFA was removed, followed by thee washes with PBS. Brains were then placed in 30% sucrose (dissolved in PBS) for 24-48 hours and subsequently frozen in cryomolds (Tissue-Tek) in O.C.T. compound (Tissue-Tek). Brains were then sectioned coronally at 35 µm on a CM3050-S cryostat (Leica). To stain, free floating sections were first blocked for 1 hour at room temperature in blocking buffer (5% goat serum, 0.3% Triton X-100) and then incubated with primary antibodies diluted in blocking buffer overnight at 4°C. The next day, sections were washed three times for 15 min per wash in PBS at room temperature. Free floating sections were then incubated in secondary antibodies diluted at 1:1000 in blocking buffer for 2 hours at room temperature. Sections were washed three times for 15 min per wash in PBS at room temperature and then mounted on Superfrost Plus microscope slides (Fisherbrand) in 10% PBS. Once the sections were dry, slides were dipped in water and allowed to dry again. Coverslips (#1.5, VWR) were affixed with DAPI Fluoromount-G (Southern Biotech). Images of the hippocampus and primary visual cortex were taken with a Nikon confocal microscope using either the 20X air objective or 60X oil objective. Imaging of immunostained sections to confirm successful viral targeting was performed on the Olympus VS200 slide scanner using a 20X air objective.

Imaging conditions were consistent between control and Astro-NL123 cKO animals. Confocal images were analyzed, while blinded, using NIS-Elements Analysis Software (Nikon).

#### Purification of Ribosome-bound mRNA using RiboTag Method

Ribosome-bound mRNA was purified using a modified version of the RiboTag method, which has been described previously (Sanz et al., 2009). Following euthanasia with isoflurane and decapitation, rapidly dissected brains were dropped in ice cold cutting solution containing the following (in mM): 228 sucrose, 2.5 KCl, 1 NaH2PO4, 26 NaHCO3, 11 glucose,7 MgSO4-7H2O, and 0.5 CaCl2. Acute coronal brain slices (300 μm) containing the dorsal hippocampus were visualized using a stereomicroscope equipped with fluorescent lamp to check for mVenus-expressed astrocytes, indicating successful viral targeting. Hippocampal area CA1 and the entire cortex region superior to CA1 from just above the alveus to the cortical service were dissected using a microknife with the aid of a stereomicroscope. CA1 and cortical tissue from a given animal was pooled, excess cutting solution was removed, and tissue was Dounce homogenized in a homogenization buffer at 10% weight/volume. Homogenate was clarified by centrifugation and 10% of the supernatant was saved as input. The remaining lysate was incubated with pre-washed anti-HA magnetic beads (Thermo) overnight at 4°C. Beads were washed 3 times with a high-salt buffer followed by elution with RLT lysis buffer with β-ME. Both input and IP samples were subjected to RNA extraction using the QIAGEN RNeasy Micro kit. RNA concentration was determined using a NanoDrop 1000 Spectrophotometer (Thermo) and stored at −80°C pending further analysis.

#### Quantitative RT-PCR (qRT-PCR)

RNA concentration was determined using a NanoDrop and equal quantities of RNA were used to synthesize cDNA with the SuperScript IV First Strand Synthesis Kit (Invitrogen, Cat# 18091050). Equal volumes of cDNA were then mixed with TaqMan Fast Virus 1-Step Master Mix (Thermo) and qRT-PCR reactions were performed using a QuantStudio 3 RT-PCR System (Thermo). We utilized PrimeTime qPCR Probe Assays (Integrated DNA Technologies), consisting of two primers and one FAM-labeled, ZEN/IBFQ-quenched 5’ nuclease probe. The Ct values for technical replicates, which were performed in duplicate for all reactions, were averaged. Data were normalized to the arithmetic mean of ActB using the 2-ΔΔCt method. For measuring the purity of cell type-specific ribosome-bound mRNA following immunoprecipitation (Figure 3C, 3F), the following assays were employed (gene, assay ID): Actb (Mm.PT.51.14022423), Aqp4 (Mm.PT.58.9080805), Sox9 (Mm.PT.58.42739087), Pdgfra (Mm.PT.56a.5639577), MBP (Mm.PT.58.28532164), P2ry12 (Mm.PT.58.43542033), and Rbfox3 or NeuN (Mm.PT.58.11398454), Nlgn1 (recognizing ex7a-ex8; Mm.PT.58.5704919), Nlgn2 (recognizing ex3-ex4; Mm.pt.58.41142345), and Nlgn3 (recognizing ex2-ex3; Mm.PT.58.31138258).

#### Slice Electrophysiology

Acute coronal brain slices (300 μm) containing the dorsal hippocampus were prepared from P45-50 Nlgn123 astrocyte conditional knockout mice and littermate controls, both of which had been injected with tamoxifen at P1. Mice were anesthetized with isoflurane and decapitated. The brain was quickly removed and dropped into into ice cold, oxygenated physiological cutting solution containing the following (in mM): 228 sucrose, 2.5 KCl, 1 NaH2PO4, 26 NaHCO3, 11 glucose,7 MgSO4-7H2O, and 0.5 CaCl2. Slices were then recovered for 30 minutes at 32°C in pre-warmed, oxygenated artificial cerebrospinal fluid (aCSF) that contained (in mM): 119 NaCl, 2.5 KCl, 1 NaH2PO4, 26 NaHCO3, 11 glucose, 2.5 CaCl2 and 1.3 MgCl2 in ddH2O. aCSF had been adjusted to a final osmolarity of 300 mOsm. Slices were then recovered at RT in oxygenated ACSF for another 60 minutes. Slices were maintained for 2-3 hours at RT in oxygenated ACSF throughout recordings. Whole-cell voltage clamp recordings were performed on CA1 pyramidal neurons while clamping the cells - 70 mV. Slices were perfused with room temperature, oxygenated ACSF at ∼2 ml / minute. Electrical signals were recorded at 5 kHz with a two channel MultiClamp 700B amplifier (Axon Instruments). They were digitalized with a Digidata 1440 digitizer (Molecular devices), which was controlled by Clampex 10.7 (Molecular Devices). Patch clamp pipettes were generated with thin-walled borosilicate glass pipettes to resistances of 3-5 MΩ. mEPSCs were recorded with an internal solution that contained (in mM): 135 cesium methanesulfonate, 8 NaCl, 10 HEPES, 2 Mg-ATP, 0.2 Na2-GTP, 0.1 spermine, 7 phosphocreatine, and 0.3 EGTA with a pH of 7.3 and adjusted to a final osmolarity of 300 mOsm. mIPSCs were recorded with an internal solution that contained (in mM): 140 mM CsCl, 2 mM MgCl2, 5 mM EGTA, 10 mM Hepes, 0.3 mM Na3-GTP, and 4 mM Na2-ATP at pH 7.35 and adjusted to a final osmolarity of 300 mOsm. For mIPSC recordings, the calcium levels were reduced to 1 mM owing to the extremely high frequency observed in CA1 pyramidal neurons. To isolate mEPSCs, recordings were performed in the presence of 0.5 μM tetrodotoxin (TTX; Cayman Chemical, Cat. 14964) and 50 μM picrotoxin (Tocris, Cat. 1128). To isolate mIPSCs, recordings were performed in the presence of 10 μM CNQX (Tocris, Cat. 1045), 50 μM D-AP5 (Tocris, Cat. 0106), and 0.5 μM TTX. Analysis was performed blindly using Clampfit 10.7 via a template search for inward currents and selected manually based on trained templates. Kinetics were calculated as 10%-90% (rise) and 90%-10% (decay) peak amplitude of currents. The first 50 events from each recordings session that was either 5 minutes long or had at least 300 events were used to generate cumulative plots.

#### Plasmids

To assess astrocyte morphology, the following plasmids were used: pAAV-GfaABC1D-Lck-mVenus and the helper plasmids pHelper and pAA5. The pAAV-GfaABC1D-Lck-mVenus construct was generated using PCR stitching to include the GfaABC1D promoter and a membrane targeting domain as described previously (Chai et al., 2017) and fused with mVenus in order to mark astrocyte membranes for analysis of astrocyte morphology. PCR stitching was also used to create a novel viral tool for capturing astrocyte-enriched ribosome-bound mRNA. The expression construct included features from previous astrocyte-targeted viral RiboTag vectors (Bravo-Ferrer et al., 2022), including the the GfaABC1D promoter, which allows efficient expression of genetic cargo in astrocytes, and RiboTag (i.e., Rpl22-HA, which allows affinity capture of ribosome-bound mRNA with HA antibodies). We also incorporated upstream of Ribotag, a membrane-targeted fluorescent reporter (i.e., Lck-mVenus) to allow visualized of targeted astrocytes, and the self-cleaving peptide, P2A, to allow separate targeting of the reporter and RiboTag. Finally, we included a 4×6T cassette of mIR-124 targeting sequences as was recently published (Gleichman et al., 2023), which significantly reduces off-target expression of genetic cargo in infected neurons.

#### AAV Preparation

For production of AAV (serotype AA5), HEK293T cells were co-transfected using the calcium phosphate method with pAAV-GfaABC1D-Lck-mVenus and the helper plasmids (pHelper and pAA5) at 4 µg per plasmid per 30 cm^2^ culture area. Approximately 12 hours after transfection, media was changed. At, ∼72 hours after the transfection, the HEK293T cells were collected using PBS with 10 mM EDTA and then spun down at 1500 x g. The cell pellets were resuspended in freezing buffer and subjected to 3 freeze-thaw cycles, alternating between 37C and a dry ice bath. Lysates were then incubated with 50 units / ml of benzonase nuclease at 37C for 30 minutes followed by a spin at 3000 x g for 30 minutes. The supernatant was loaded into discontinuous iodixanol gradient and centrifuged at 65,000 rpm for 3 hours. The 40% iodixanol fraction was collected and concentrated using 100,000 MWCO centricon columns. Filtrate was washed with several charges of MEM and then aliquoted and stored at −80C until use. Although viral titer was not determined, purified AAV was injected in vivo at various dilutions in order to identify a dilution that allowed sparse targeting of astrocytes for morphological reconstruction; thus, final viral titer was relatively low.

AAV2/5-GfaABC1D-Lck-mVenus-P2A-Rpl22-HA-4×6T was produced by the Stanford Gene Vector and Virus Core (RRID:SCR_023250). The titer was determined to be 2.10E+13 and 3.00E+13 using assays that recognize the WPRE and ITR sequences, respectively.

#### Stereotactic Injections

Prior to surgery, P21 mice were anesthetized with tribromoethanol (Avertin) at a dosage of 125-300 mg/kg and Buprenorphine Sustained Release was administered at 0.5 mg/kg to provide analgesia. Mouse heads were shaved and then cleaned with Betadine prior to incision of the scalp. Using a stereotactic rig (Kopf) for targeting, AAV expressing membrane-targeted mVenus under the control of the GFAP promoter was injected into CA1 (coordinates from bregma: A-P −1.8, M-L ±1.15, D-V −1.4) or V1 (coordinates from bregma: A-P −3.99, M-L ±2.6, D-V - 1.5). Virus was injected through a glass pipette attached to a syringe pump (SP101i, World Precision Instruments) at a speed of 0.15 µl/min at a volume that was empirically determined to sparsely infect astrocytes (0.5 µl). Following injection, the incision was closed with 4-0 nylon sutures (Unify). Mice were allowed to recover in a clean cage on a heating pad prior to transfer to a clean home cage. Sutures were removed 10 days post-surgery.

For validating Nlgn123 knockout efficiency, P35-P40 mice were injected with AAV2/5-GfaABC1D-Lck-mVenus-P2A-Rpl22-HA-4×6T diluted at 1:1 in DMEM (final titer is 1.05E+13 and 1.5E+13 based on WPRE and ITR sequences, respectively). 0.45 μl per hemisphere (coordinates from bregma: A-P −1.8, ML ± 1.45, D-V −1.15 to −1.45 with 0.15 μl per 0.15 step).

#### Astrocyte Morphology

Mice were stereotactically injected with AAV expressing membrane-targeted mVenus under the control of the GFAP promoter at P21 and then perfused, as described in the immunohistochemistry section, at P35. Brains were post-fixed in 4% PFA for 24 hours at 4°C. Brains were then washed 3 times in PBS and then vibratome sectioned at 100µm in PBS. Sections were counterstained with DAPI (D8417, Sigma) and mounted on gelatin-coated slides (FD Neurotechenologies, Cat# PO101) in 10% PBS. Once sections were dry, slides were dipped in water and again allowed to dry. Coverslips (#1.5, VWR) were affixed with Fluoromount-G (Southern Biotech). Images were taken on a Zeiss AiryScan microscope using the same imaging parameters for controls and Astro-Nlgn123 cKO animal. While blinded, astrocyte morphologies were reconstructed with 3D rendering in Imaris and astrocyte volumes were measured.

#### Data Analysis and Statistics

For electrophysiology experiments, data were analyzed with Clampfit 10.7 (Molecular Devices). For immunoblot, immunohistochemistry, and astrocyte morphology experiments, unpaired two-tailed t-tests were used to assess statistical significance. For electrophysiology experiments, unpaired two-tailed t-tests were used to analyze data plotted in bar graphs (e.g., rise time) and Kolmogorov-Smirnov tests were used to assess statistical significance of cumulative curves. For bar graphs, data are depicted as means ± SEM. For all experiments, significance is indicated by * p<0.05, **p<0.01 or ****p<0.0001.

## DATA AVAILABILITY

All raw data for this study are deposited in the Stanford Digital Repository (https://purl.stanford.edu/ys873rd9462) and are freely available.

## ACKNOWLEDGEMENTS

We thank Dr. Yi Han Ng for her enormous help in submitting the voluminous raw data for this paper to the Stanford Digital Repository. This study was supported by grants from the NIMH (K01-MH123788 to J.H.T. and R01MH092931 to T.C.S.) and the BBRF J.H.T. and facilitated by the Stanford University Cell Sciences Imaging Core Facility (RRID:SCR_017787) and the Stanford Gene Vector and Virus Core (RRID:SCR_023250).

## AUTHOR CONTRIBUTIONS

S.G., J.H.T., and G.N. performed all experiments except for the experiments with human neurons that were performed by J.W.; S.G., J.H.T. and T.C.S. designed the experiments, and all authors analyzed the data and wrote the paper.

## SUPPLEMENTARY FIGURES and FIGURE LEGENDS

**Figure S1:**
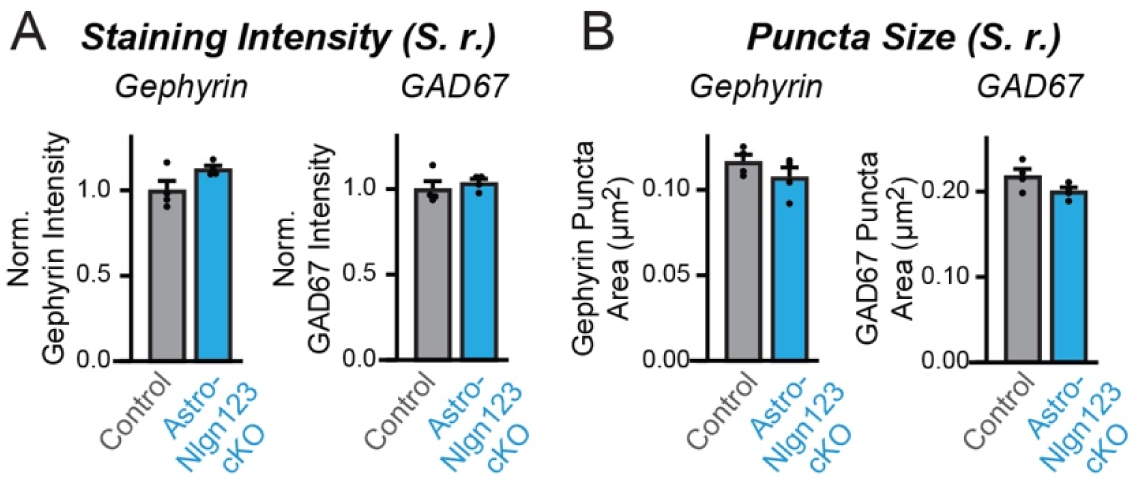
Conditional deletion of astrocytic *Nlgn1-3* at P1 does not alter the number or size of inhibitory synapses in the CA1-region *Stratum Radiatum*. (**A**) Quantification of total Gephyrin (left) or GAD67 (right) immunofluorescence in CA1 *Str. Radiatum* from astrocyte Nlgn1-3 cKO and littermate control mice. Images of hippocampal sections were taken at 60X magnification. Total immunofluorescent signal was first internally normalized to MAP2 and then to average gephyrin (left) or GAD67 (right) immunofluorescence level in control mice. (**B**) Quantification of puncta density for Gephyrin (left) and GAD67 (right) CA1 *Str. Radiatum* from astrocyte Nlgn1-3 cKO and littermate control mice. Representative images are shown in Fig. 4G. Data are means ± SEM with statistical significance determined by unpaired two-tailed t-test (n=4, 2 male & 2 female).

**Figure S2:**
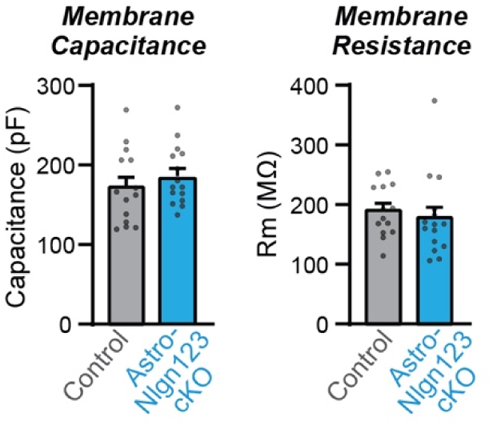
Conditional deletion of astrocytic *Nlgn1-3* at P1 does not alter CA1 pyramidal neuron membrane properties. (**A**) Summary graph of membrane capacitance from CA1 pyramidal neurons in acute slices from astrocyte Nlgn1-3 cKO and littermate controls injected with tamoxifen at P1 and recorded at P44 – P50. (**B**) Same as (**A**) but for membrane resistance. Data are means ± SEM with statistical significance determined by unpaired two-tailed t-test (n = 14-15 cells / 3 mice per genotype).

**Figure S3:**
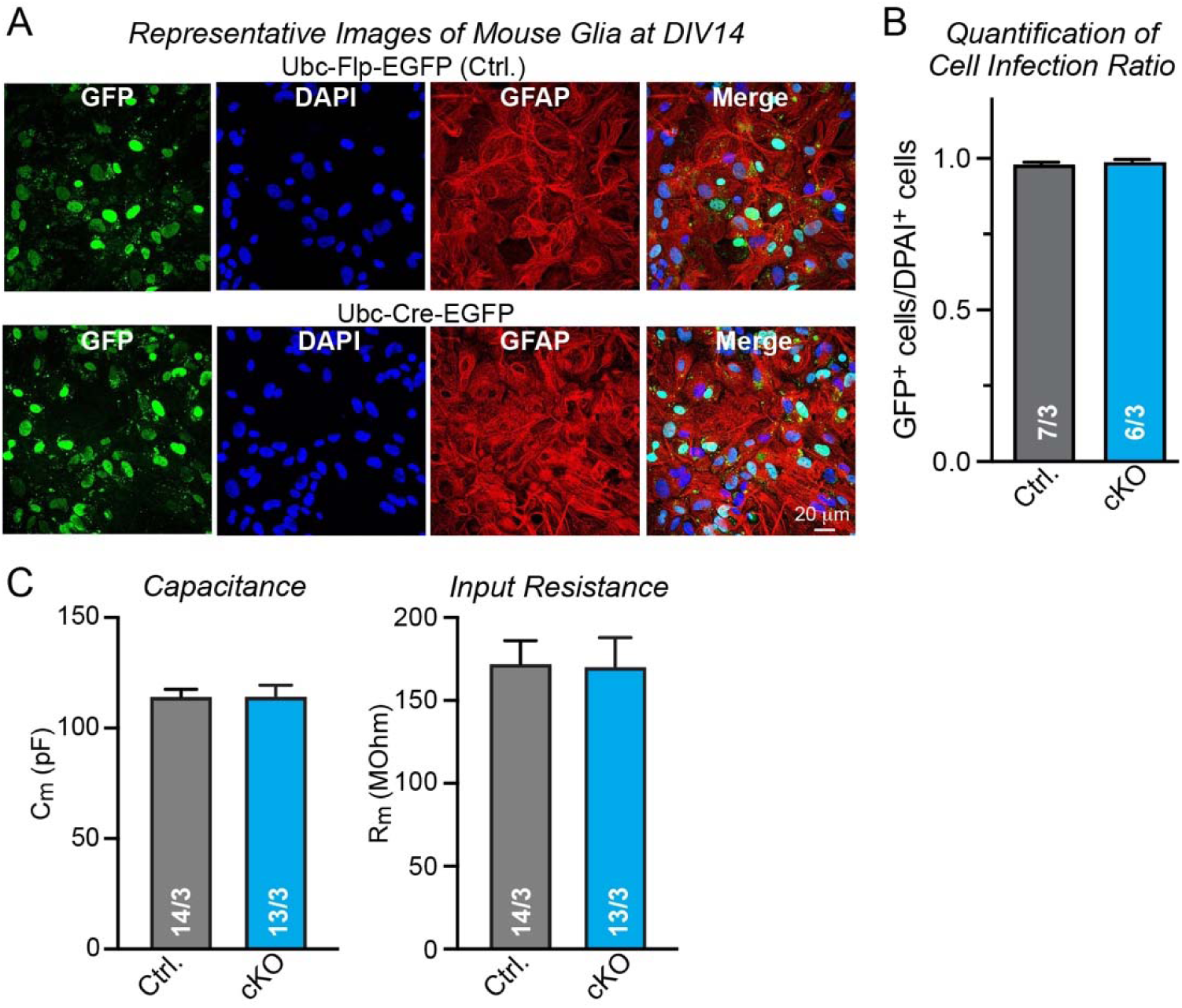
Further data characterizing the effect of a deletion of glial *Nlgn1-3* on co-cultured human neurons. (**A**) Representative images of primary mouse glia cultures from *Nlgn1-4* quadruple cKO mice infected with lentiviruses encoding FLP-EGFP (Ctrl.) or Cre-EGFP fusion proteins and stained for GFP, DAPI, and GFAP as glial marker. Note that the levels of EGFP expression differ among glia cells but that based on qRT-PCR measurements the lower levels of Cre-EGFP in some glial cells are sufficient for complete recombination of *Nlgn1*, *Nlgn2*, and *Nlgn3* genes (*Nlgn4* is constitutively deleted). (**B**) Summary graph of the ratio of EGFP-expressing to DAPI-stained cells demonstrates that nearly all cells in the culture are infected by the lentiviruses. (**C**) Summary graphs of the membrane capacitance (left) and input resistance (right) monitored in human neurons that are co-cultured with mouse glia expressing or lacking all neuroligins (Nlgn1-4). Data are means ± SEM with statistical significance determined by unpaired two-tailed t-test (n = 14-15 cells / 3 mice per genotype).

